# Local genomic estimates provide a powerful framework for haplotype discovery

**DOI:** 10.1101/2025.08.28.672830

**Authors:** Will Shaffer, Victor Papin, Seema Yadav, Kai P. Voss-Fels, Lee T. Hickey, Ben J. Hayes, Eric G. Dinglasan

**Affiliations:** Queensland Alliance for Agriculture and Food Innovation, The University of Queensland, Brisbane, Queensland, 4072, Australia; Institute of Plant Breeding, Hochschule Geisenheim University, 65366 Geisenheim, Germany

**Keywords:** local GEBV, linkage disequilibrium, GWAS, genomic selection, haplotype, haploblock

## Abstract

Quantitative trait loci (QTL) discovery studies on diversity panels or breeding populations typically use genome-wide association studies (GWAS) to estimate marker effects. For plant and animal breeding applications, researchers increasingly recognize the potential benefits of identifying superior haplotypes (markers in linkage disequilibrium; LD) rather than relying on single markers, as traditional approaches inefficiently account for cumulative signals from incomplete LD with QTL or split effects when multiple markers are in high LD with QTL. Using the genomic prediction framework, the local genomic estimated breeding values (localGEBV) method was developed in animal breeding and has been adopted in crop haplotype mapping studies; however, no study has thoroughly quantified the utility of this method or systematically compared outcomes to traditional GWAS approaches. Here, we characterized a strategy to group markers in chromosomal segments based on LD (haplotype blocks or haploblocks), computed localGEBV as a linear contrast of marker effects within each haploblock, and utilised the variance of localGEBV to enhance QTL discovery compared to traditional GWAS. Marker effects for localGEBV were estimated with ridge-regression best linear unbiased prediction (rrBLUP) and BayesR, with results compared to two common GWAS approaches. Using the barley row-type trait, we demonstrated that localGEBV improved QTL discovery and phenotypic prediction compared to single markers. Furthermore, localGEBV results were robust to the choice of prior marker assumptions and blocking parameters, enabling flexibility in fine or broad-scale QTL mapping. Overall, our findings establish localGEBV as a haplotype-based strategy capable of leveraging localized genomic effects to improve QTL discovery and, potentially, genomic selection.

## Introduction

Understanding the genetic basis of trait variation is fundamental to advancing crop and livestock improvement programs. Identifying quantitative trait loci (QTL) or genomic regions harbouring mutations that contribute to trait variation provides crucial insights into genetic architecture and enables targeted breeding strategies. Traditional genome-wide association studies (GWAS) detect QTL by testing associations between individual genetic markers and phenotypes using linear regression, while modern approaches utilize linear mixed models, Bayesian hierarchical models, or multi-stage methods. A significant association indicates the marker is in linkage disequilibrium (LD) with a causal mutation.

The single-marker approach proves effective in populations with limited LD, such as human populations with large effective population sizes. However, this approach lacks precision in populations characterized by more extensive LD due to small effective population sizes – a common feature of crop and livestock populations. In these populations, markers that are in LD often split the QTL effect when estimated simultaneously, and individual markers may capture only fragments of the total QTL effect through imperfect LD relationships (Meuwissen and Goddard, 2000; Marchini et al., 2004; Goddard, 2009; Zhang et al., 2010). This dilution of signal across multiple correlated markers reduces the power to detect causal variants.

To address these limitations, chromosome segments in LD can be leveraged through haplotype-based approaches that capture the collective signal across multiple linked markers (Meuwissen and Goddard, 2000; The Bovine HapMap Consortium et al., 2009; Comadran et al., 2011; Joukhadar et al., 2017). These methods group markers into haplotype blocks, typically defined as LD-based chromosomal segments (LD blocks or haploblocks), and analyse the combined effects within each block. One promising haplotype-based method utilizes the variance of local genomic estimated breeding values (localGEBV), which represents the variance of haplotype effects within a block. This approach has demonstrated the ability to detect QTL where causal mutations are known in livestock populations (Fan et al., 2011; Kemper et al., 2015; Van Den Berg et al., 2019; Xiang et al., 2021). Unlike traditional GWAS that tests markers individually, the localGEBV approach, derived from genomic prediction methods, begins with fitting all single nucleotide polymorphism (SNP) simultaneously (Meuwissen et al., 2001), counteracting shrinkage effects and the dilution of marker effects when multiple markers tag the same QTL. Theoretical advantages of the localGEBV approach include reduced susceptibility to false positives from population stratification, less severe over-estimation of effects, and improved QTL detection power, all stemming from the simultaneous fitting of all SNP rather than individual testing (Yang et al., 2014).

The localGEBV method has also shown promise in crop applications. Voss-Fels et al. (2019) first applied this approach in wheat to identify beneficial haplotypes for yield, though their focus was on breeding program improvements rather than QTL discovery methodology. Others have applied the same method to identify haplotypes important for disease resistance in barley and wheat (Tong et al., 2024; Roy et al., 2025; Vo Van-Zivkovic et al., 2025), stay-green and root traits in barley (Brunner et al., 2024; Alahmad et al., 2025; Aldiss et al., 2025), and chickpea seed weight, flowering time, and plant height (Varshney et al., 2021; Akinlade et al., 2025).

Despite these promising developments, critical knowledge gaps limit our understanding and application of haplotype-based QTL detection. No direct comparison between localGEBV and standard GWAS approaches has been conducted to empirically evaluate their relative performance. Previous applications in QTL discovery in livestock have relied on sliding window approaches, whereas in crop species, haplotype discovery focused on biologically meaningful LD-based blocks. However, the effect of fundamental methodological aspects, including optimal marker effect priors and blocking parameters, remain poorly characterized for QTL discovery.

Here, we address these gaps through comprehensive evaluation of the localGEBV approach using a barley global diversity panel genotyped with 40K SNP, focusing on spike row morphology, a trait with complex genetic architecture distinguishing 2-row from 6-row varieties. We construct LD-based haploblocks, investigate localGEBV estimation using ridge regression best linear unbiased prediction (rrBLUP) and BayesR to account for different marker effect priors (Meuwissen et al., 2001; Erbe et al., 2012), and directly compare QTL detection capabilities against established multi-locus GWAS methods (FarmCPU and BLINK). Our findings demonstrate that haplotype-based localGEBV offers superior precision and power for QTL discovery in populations with extensive LD by detecting multiple known QTL compared to modern GWAS methods, providing a robust framework applicable across plant and animal breeding programs.

## Results

### Barley accessions display distinct genetic differentiation for row-type phenotypes

Principal component analysis revealed substantial genetic differentiation among the barley accessions, with PC1 and PC2 explaining 30.4% and 14.4% of variation, respectively. *K-means* clustering identified three main genetic clusters (Figure 1A): Cluster 1 (313) comprised predominantly 2-row accessions, while Clusters 2 (297) and 3 (180) were primarily composed of 6-row accessions (Figure 1B and 1D).

**Figure 1.**
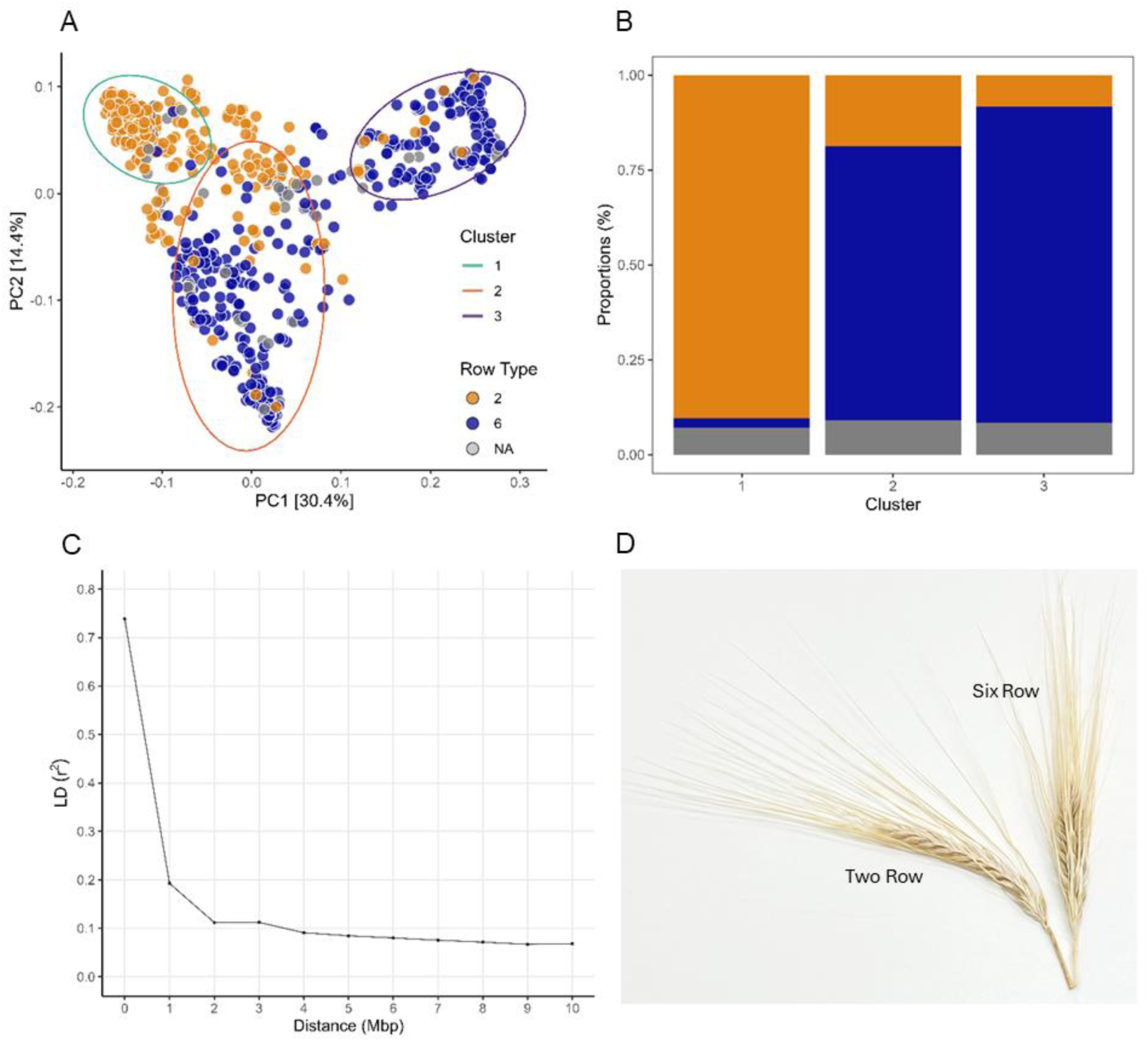
Population structure and genetic relationships of barley germplasm. (**A**) Principal component analysis of *N* = 790 barley accessions using 9,050 XT SNP chip markers, revealing three distinct genetic clusters. Shown are PCs 1 and 2 from Roger’s genetic distance matrix, accounting for 44.8% cumulative explained variation. The optimum number of clusters were *clusters* = 3: Cluster 1 (313), Cluster 2 (297), and Cluster 3 (180). *Colours* represent the row spike phenotype i.e. 2-row is *orange*, 6-row is *blue*. (**B**) Proportions of barley accessions across the three clusters: Cluster 1 (2-row = 283, 6-row = 8, NA = 22), Cluster 2 (2-row = 56, 6-row = 214, NA = 27), Cluster 3 (2-row = 15, 6-row = 150, NA = 15), where NA represents accessions with ambiguous information and ‘irregular’ type. (**C**) Genome-wide LD decay as a function of physical distance (Mbp). LD decay is shown at an average distance (Mbp) between adjacent SNP based on the XT SNP chip physical position using cv Morex assembly v2 and visualised up to 10 Mbp. (**D**) An example of the two-row and six-row phenotypes.

LD analysis demonstrated high initial LD (𝑟^2^ = 0.74) within distances <0.1 Mbp, followed by rapid decay to 𝑟^2^ = 0.19 at 1 Mbp, with subsequent gradual decay thereafter (Figure 1C).

### LD-based haploblock construction reveals diverse genomic segmentation patterns

The construction of chromosome segments using varying LD parameters generated distinct haploblock configurations. Relaxed parameters (lower 𝑟^2^ threshold and higher marker tolerance, 𝑡𝑜𝑙, which allowed some markers in the block to not meet the LD threshold) generated fewer haploblocks total with more markers per haploblock, while stringent parameters (higher 𝑟^2^threshold, lower marker tolerance) produced more numerous haploblocks that were predominantly single-marker blocks (Supplementary Figure 1). For instance, at 𝑟^2^ = 0.1 with zero marker tolerance, 3,738 haploblocks (447,691 possible haplotypes) were constructed, with 60.27% comprising single SNP markers. Increasing the 𝑟^2^threshold to 0.5 resulted in 6,781 haploblocks (103,857 possible haplotypes), where 73.72% consisted of single SNP markers.

Increasing marker tolerance substantially reduced haploblock count while expanding block size and possible haplotype numbers. At 𝑟^2^ = 0.1 with marker 𝑡𝑜𝑙 = 3, only 1,096 haploblocks were constructed (2,890,043 possible haplotypes). The largest observed haploblock (364 SNP markers spanning 299 Mbp on chromosome 5H) contained 374,192 possible haplotypes.

### Validation of localGEBV approach for mapping *VRS1* gene in chromosome 2H

To determine whether haploblock variance was influenced by segment size, we applied a generalized additive linear regression model, which revealed no meaningful linear relationship between haploblock variance and block size (number of markers; 𝑅^2^ = 0.07, *P* < 2e-16; Figure 2, Supplementary Figure 3). Instead, a significant and meaningful relationship was observed between haploblock variance and haplotype effects (𝑅^2^ = 0.48, *P* < 2e-16).

**Figure 2.**
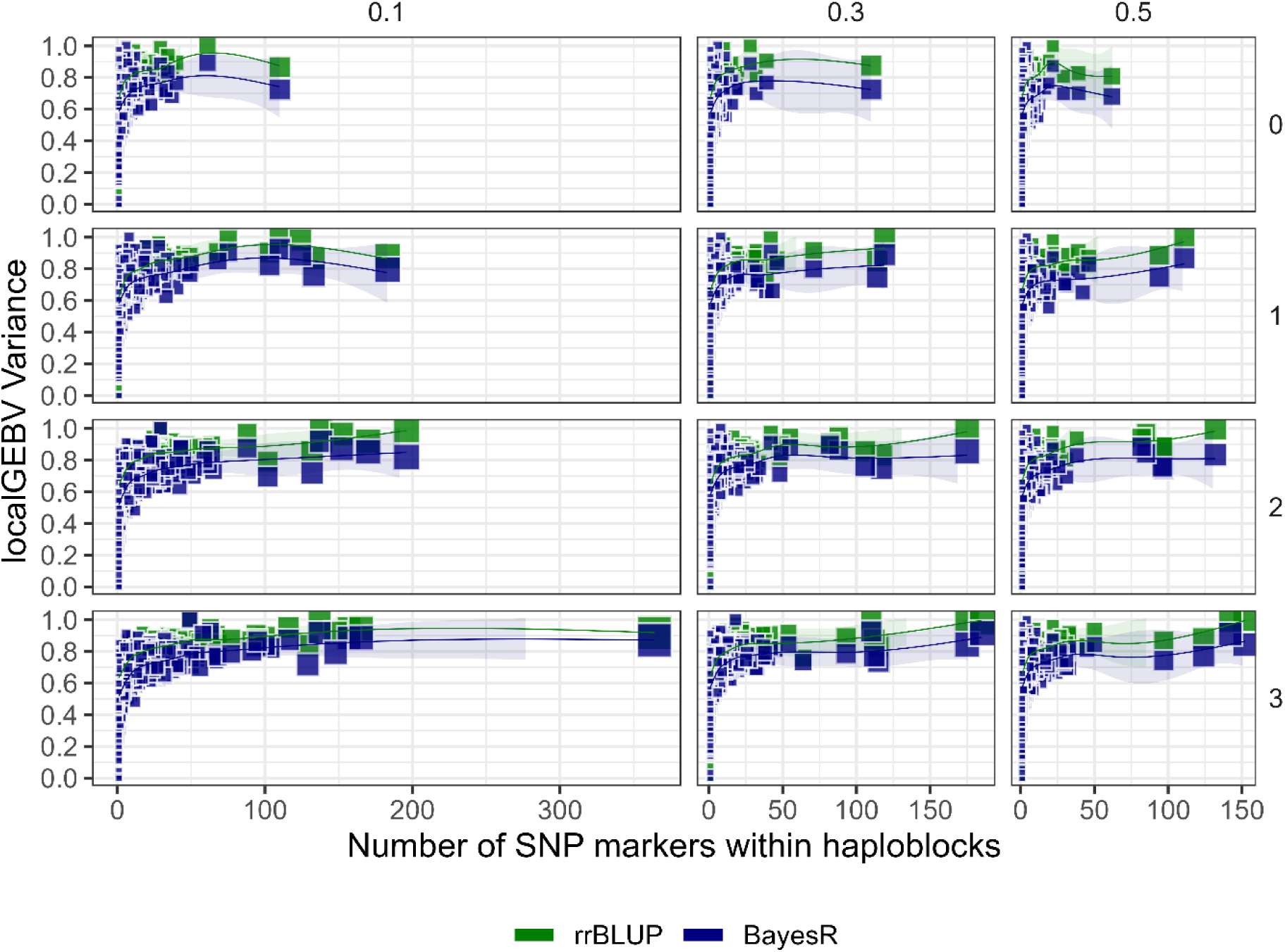
Relationship between haploblock variance and haploblock size. Haploblock variance derived from localGEBV estimates using rrBLUP (*green*) and BayesR (*blue*), plotted on the *y-axis*. Block size is defined by the number of SNP markers within haploblocks, plotted on the *x-axis*. In the figure, different LD parameters are shown as facets: *column facets* represent LD thresholds of 𝑟^2^ ∈ {0.1, 0.3, 0.5}; and *row facet* as marker tolerance, 𝑡𝑜𝑙 ∈ {0, 1, 2, 3}. The localGEBV variance was scaled using a min-max scaling of the log_10_transformed variance. Blocks were connected using a generalized additive model for non-linear relationships to represent trends between haploblock size and localGEBV variance.

We validated the localGEBV approach by examining the well-characterized *VRS1* gene on chromosome 2H, which determines row-type phenotype (Komatsuda et al., 2007). Marker effect estimation using rrBLUP showed relatively small effects distributed across the chromosome, with modest increase in the *VRS1* region (Supplementary Figure 2). In contrast, BayesR attributed larger effects to SNP collocated with *VRS1* compared to surrounding markers. Analysis of localGEBV variance across different LD parameters consistently identified haploblocks peaking at the *VRS1* region (Figure 3A). Both rrBLUP and BayesR detected maximum variance at 𝑟^2^ = 0.1 and marker tolerance, 𝑡𝑜𝑙 = 3, co-locating with *VRS1* and with localGEBV variance of 0.0081 and 0.0133, respectively (Figure 3A; Table 1).

**Figure 3.**
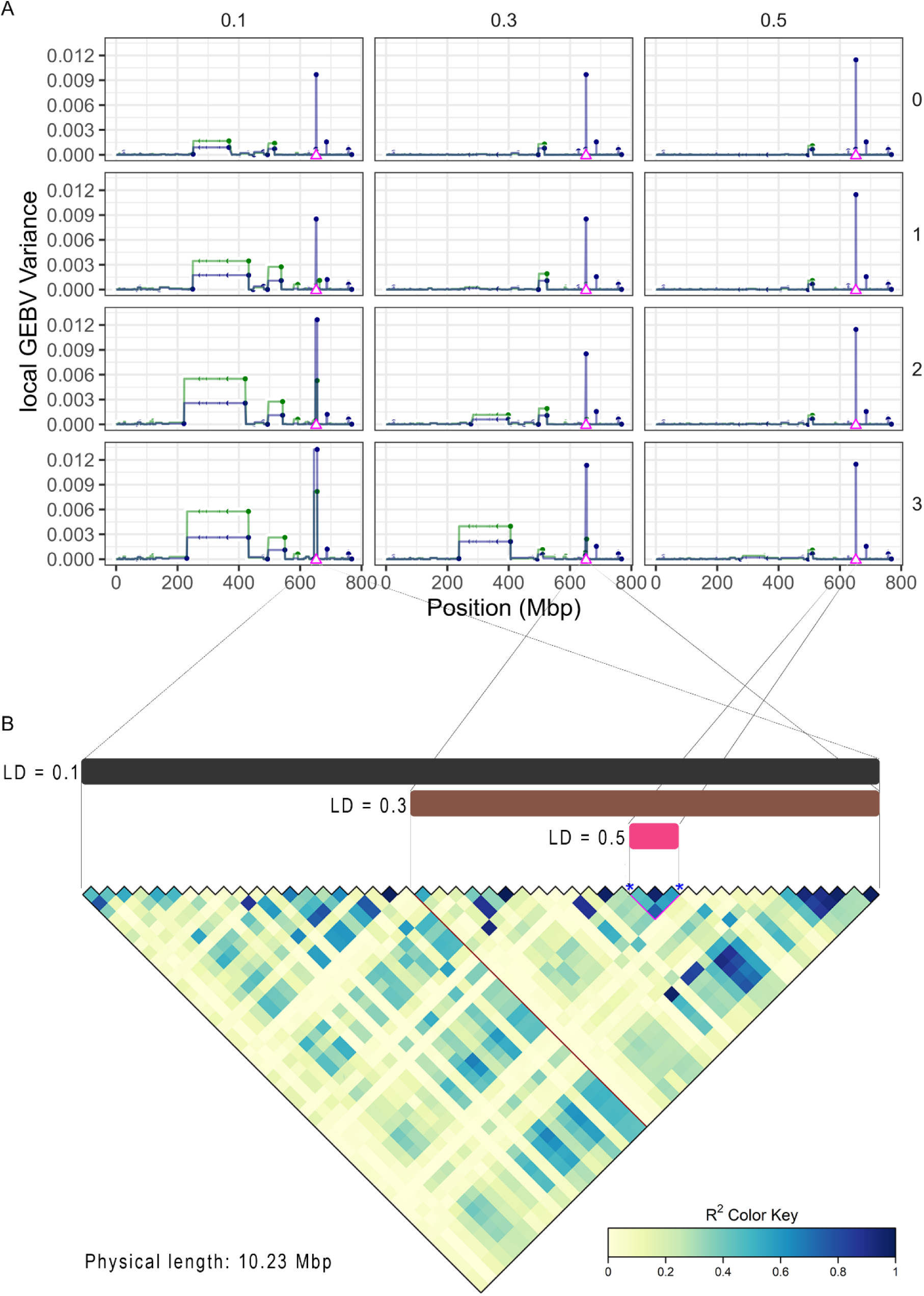
LocalGEBV variance of haploblocks in chromosome 2H. The *VRS1* gene (marked as Δ) is co-located at 652.03 Mbp based on the XT SNP chip physical position using cv Morex assembly v2. (**A**) The localGEBV variance profiles across chromosome 2H estimated using rrBLUP (*green*) or BayesR (*blue*). Haploblocks are constructed according to different LD parameters: where *column facet* represents LD threshold at 𝑟^2^ ∈ {0.1, 0.3, 0.5} ; and *row facet* as 𝑡𝑜𝑙 ∈ {0, 1, 2, 3}. (**B**) Varying sizes of the haploblocks based on LD threshold show fine-scale mapping in the *VRS1* region. Haploblocks constructed at 𝑟^2^ ∈ {0.1, 0.3, 0.5} with 𝑡𝑜𝑙 ∈ {0, 1, 2, 3}. Shown together is the corresponding LD heatmap.

**Table 1.**
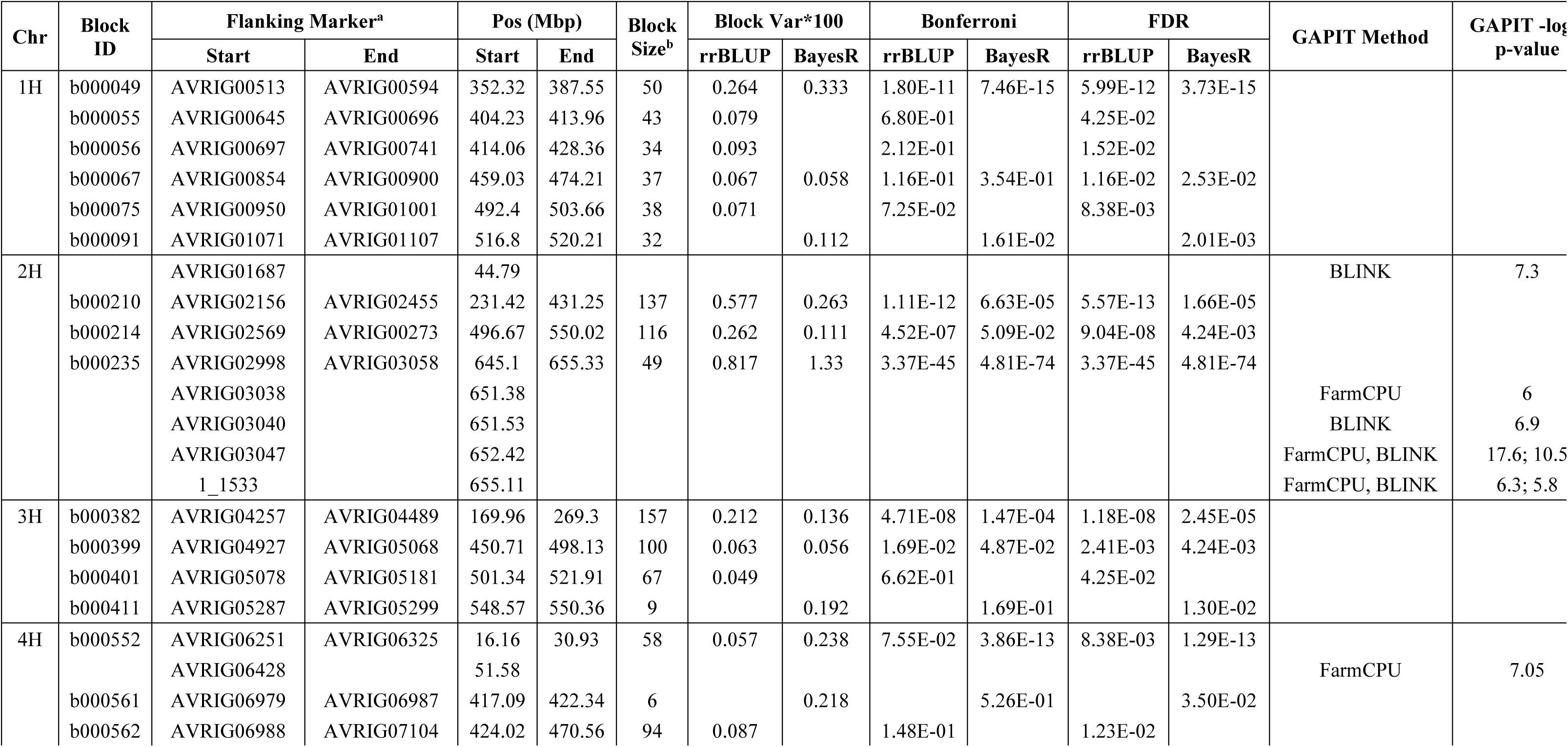
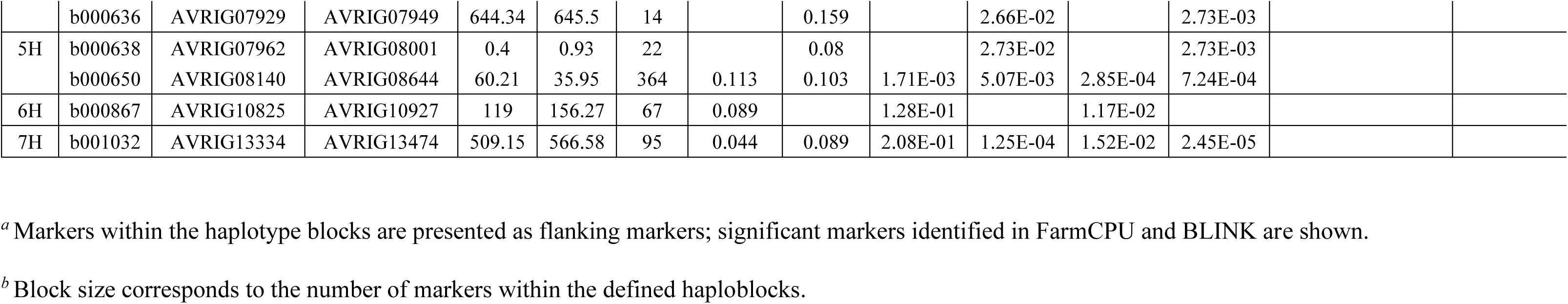
Summary statistics of high variance haploblocks for row-spike phenotype from localGEBV estimated using rrBLUP and BayesR, and significant markers identified using FarmCPU and BLINK. Only haploblocks with localGEBV Bonferroni or False Discovery Rate (FDR) corrected p-values are displayed.

| Chr | Block ID | Flanking Marker<sup>a</sup> |  | Pos (Mbp) |  | Block Size<sup>b</sup> | Block Var*100 |  | Bonferroni |  | FDR |  | GAPIT Method | GAPIT -log p-value |
| --- | --- | --- | --- | --- | --- | --- | --- | --- | --- | --- | --- | --- | --- | --- |
|  |  | Start | End | Start | End |  | rrBLUP | BayesR | rrBLUP | BayesR | rrBLUP | BayesR |  |  |
| 1H | b000049 | AVRIG00513 | AVRIG00594 | 352.32 | 387.55 | 50 | 0.264 | 0.333 | 1.80E-11 | 7.46E-15 | 5.99E-12 | 3.73E-15 |  |  |
|  | b000055 | AVRIG00645 | AVRIG00696 | 404.23 | 413.96 | 43 | 0.079 |  | 6.80E-01 |  | 4.25E-02 |  |  |  |
|  | b000056 | AVRIG00697 | AVRIG00741 | 414.06 | 428.36 | 34 | 0.093 |  | 2.12E-01 |  | 1.52E-02 |  |  |  |
|  | b000067 | AVRIG00854 | AVRIG00900 | 459.03 | 474.21 | 37 | 0.067 | 0.058 | 1.16E-01 | 3.54E-01 | 1.16E-02 | 2.53E-02 |  |  |
|  | b000075 | AVRIG00950 | AVRIG01001 | 492.4 | 503.66 | 38 | 0.071 |  | 7.25E-02 |  | 8.38E-03 |  |  |  |
|  | b000091 | AVRIG01071 | AVRIG01107 | 516.8 | 520.21 | 32 |  | 0.112 |  | 1.61E-02 |  | 2.01E-03 |  |  |
| 2H |  | AVRIG01687 |  | 44.79 |  |  |  |  |  |  |  |  | BLINK | 7.3 |
|  | b000210 | AVRIG02156 | AVRIG02455 | 231.42 | 431.25 | 137 | 0.577 | 0.263 | 1.11E-12 | 6.63E-05 | 5.57E-13 | 1.66E-05 |  |  |
|  | b000214 | AVRIG02569 | AVRIG00273 | 496.67 | 550.02 | 116 | 0.262 | 0.111 | 4.52E-07 | 5.09E-02 | 9.04E-08 | 4.24E-03 |  |  |
|  | b000235 | AVRIG02998 | AVRIG03058 | 645.1 | 655.33 | 49 | 0.817 | 1.33 | 3.37E-45 | 4.81E-74 | 3.37E-45 | 4.81E-74 |  |  |
|  |  | AVRIG03038 |  | 651.38 |  |  |  |  |  |  |  |  | FarmCPU | 6 |
|  |  | AVRIG03040 |  | 651.53 |  |  |  |  |  |  |  |  | BLINK | 6.9 |
|  |  | AVRIG03047 |  | 652.42 |  |  |  |  |  |  |  |  | FarmCPU, BLINK | 17.6; 10.5 |
|  |  | 1_1533 |  | 655.11 |  |  |  |  |  |  |  |  | FarmCPU, BLINK | 6.3; 5.8 |
| 3H | b000382 | AVRIG04257 | AVRIG04489 | 169.96 | 269.3 | 157 | 0.212 | 0.136 | 4.71E-08 | 1.47E-04 | 1.18E-08 | 2.45E-05 |  |  |
|  | b000399 | AVRIG04927 | AVRIG05068 | 450.71 | 498.13 | 100 | 0.063 | 0.056 | 1.69E-02 | 4.87E-02 | 2.41E-03 | 4.24E-03 |  |  |
|  | b000401 | AVRIG05078 | AVRIG05181 | 501.34 | 521.91 | 67 | 0.049 |  | 6.62E-01 |  | 4.25E-02 |  |  |  |
|  | b000411 | AVRIG05287 | AVRIG05299 | 548.57 | 550.36 | 9 |  | 0.192 |  | 1.69E-01 |  | 1.30E-02 |  |  |
| 4H | b000552 | AVRIG06251 | AVRIG06325 | 16.16 | 30.93 | 58 | 0.057 | 0.238 | 7.55E-02 | 3.86E-13 | 8.38E-03 | 1.29E-13 |  |  |
|  |  | AVRIG06428 |  | 51.58 |  |  |  |  |  |  |  |  | FarmCPU | 7.05 |
|  | b000561 | AVRIG06979 | AVRIG06987 | 417.09 | 422.34 | 6 |  | 0.218 |  | 5.26E-01 |  | 3.50E-02 |  |  |
|  | b000562 | AVRIG06988 | AVRIG07104 | 424.02 | 470.56 | 94 | 0.087 |  | 1.48E-01 |  | 1.23E-02 |  |  |  |
|  | b000636 | AVRIG07929 | AVRIG07949 | 644.34 | 645.5 | 14 |  | 0.159 |  | 2.66E-02 |  | 2.73E-03 |  |  |
| 5H | b000638 | AVRIG07962 | AVRIG08001 | 0.4 | 0.93 | 22 |  | 0.08 |  | 2.73E-02 |  | 2.73E-03 |  |  |
|  | b000650 | AVRIG08140 | AVRIG08644 | 60.21 | 35.95 | 364 | 0.113 | 0.103 | 1.71E-03 | 5.07E-03 | 2.85E-04 | 7.24E-04 |  |  |
| 6H | b000867 | AVRIG10825 | AVRIG10927 | 119 | 156.27 | 67 | 0.089 |  | 1.28E-01 |  | 1.17E-02 |  |  |  |
| 7H | b001032 | AVRIG13334 | AVRIG13474 | 509.15 | 566.58 | 95 | 0.044 | 0.089 | 2.08E-01 | 1.25E-04 | 1.52E-02 | 2.45E-05 |  |  |

<sup>a</sup> Markers within the haplotype blocks are presented as flanking markers; significant markers identified in FarmCPU and BLINK are shown.
<sup>b</sup> Block size corresponds to the number of markers within the defined haploblocks.

Closer examination of the highest variance haploblock in chromosome 2H identified progressive refinement from 49 SNP spanning 10.23 Mbp at 𝑟^2^ = 0.1 to 4 SNP spanning 0.88 Mbp at 𝑟^2^ = 0.5 (Figure 3B), denoting the trade-off between flexibility of block discovery power and fine-mapping resolution with choice in blocking parameters. BayesR consistently produced higher variance estimates at the *VRS1* region compared to rrBLUP, likely due to its heterogeneous variance assumptions allowing larger effects for SNP in high LD with QTL. Additionally, rrBLUP identified dispersed regions of additional high variance haploblocks in chromosome 2H, particularly within 231.42 to 431.25 Mbp when using relaxed LD parameters. These regions diminished in BayesR at higher LD thresholds, consistent with its assumption that most SNP have no effect on the trait (Supplementary Figure 2).

### Ability of localGEBV to detect QTL

We first assessed QTL detection using the mean haploblock variance and the blocking parameters that maximised the haploblock variance at the *VRS1* locus, which corresponded to blocking parameters of 𝑟^2^ = 0.1 and 𝑡𝑜𝑙 = 3. With these parameters, we identified 𝑟𝑟𝐵𝐿𝑈𝑃 = 158 and 𝐵𝑎𝑦𝑒𝑠𝑅 = 136 high-variance haploblocks that surpassed the mean haploblock variance from the total of 𝐽 = 1096 haploblocks (Figure 6), of which 96 were in common.

We then employed a chi-squared distribution to generate Bonferroni and False Discovery Rate (FDR) probability values (Table 1). At an FDR critical value of 0.05, 𝑟𝑟𝐵𝐿𝑈𝑃 = 16 and 𝐵𝑎𝑦𝑒𝑠𝑅 = 15 haploblocks were identified as significant. Comparative GWAS analysis using FarmCPU and BLINK identified 6 significant markers: 5 on chromosome 2H (2 detected by both methods) and 1 on 4H (FarmCPU only). Compared to the GWAS results, localGEBV identified 21 unique high variance haploblocks distributed across all chromosomes, with 10 detected by both rrBLUP and BayesR, and 6 and 5 uniquely identified by each method, respectively (Table 1, Figure 4).

**Figure 4.**
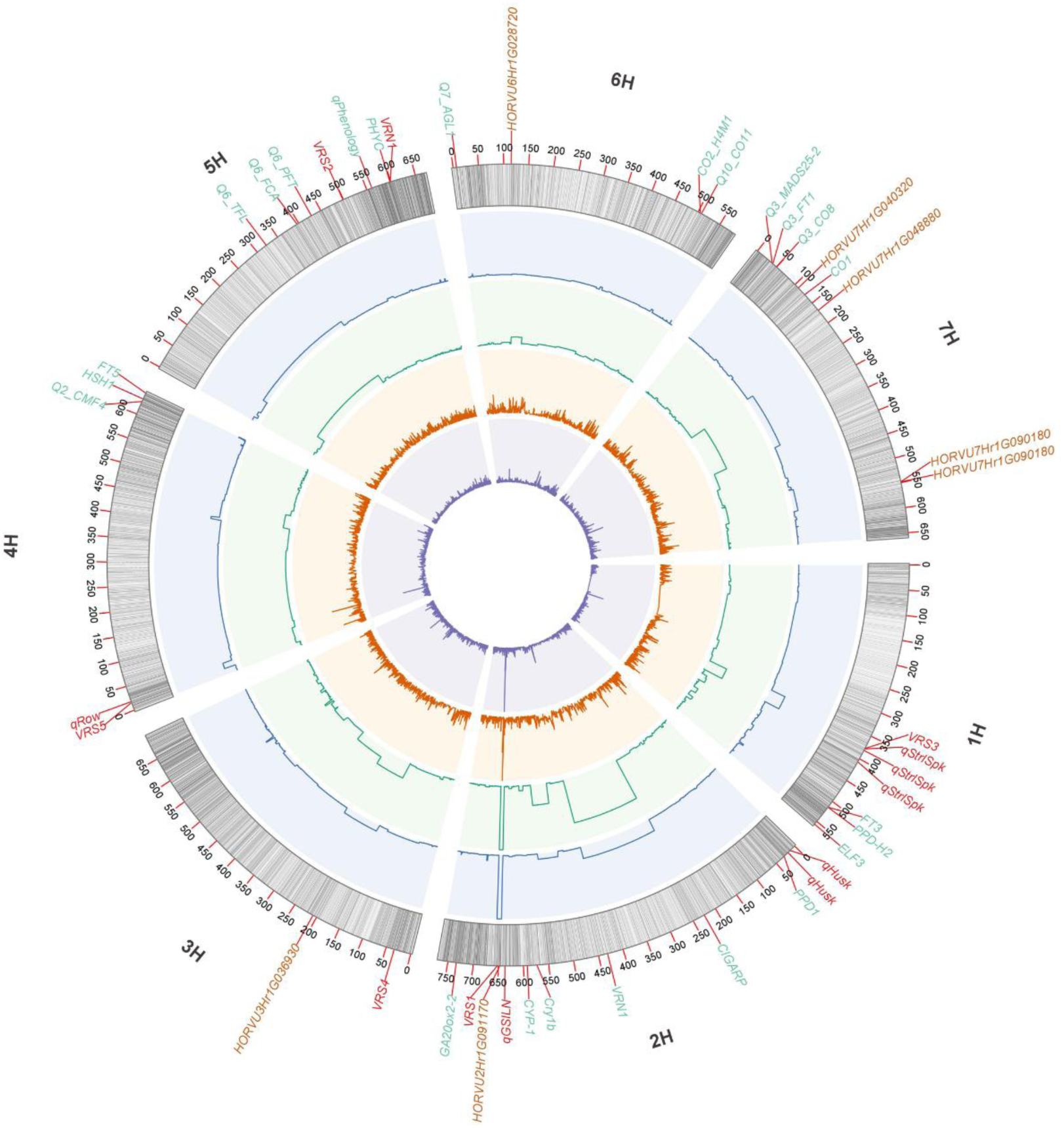
Detection of haploblocks containing QTL. Physical position in barley chromosome of SNP markers associated with row-type spike architecture detected from localGEBV and GWAS**. *Tracks 1 to 5 (outer to innermost):* Track 1** Barley chromosome showing physical position (Mbp) based on cv Morex assembly v2 from the 40K XT SNP chip. SNP markers are coloured in *grey*. Colocation of known genes and QTL from the literatures is shown: row-type known genes (*VRS3* in 1H, *VRS1* in 2H, *VRS4* in 3H, *VRS5* in 4H, *VRS2* in 5H; *dark red*), phenology genes (*FT3*, *PPD1*, *CIGARP*, *Cry1b*, *CYP-1*, *GA20ox2*-2, Q2_*CMF4*, *HSH1*, *FT5*, Q6_*TFL*, Q6_*FCA*, *Q6_PFT*, *PHYC*, *Q7_AGL1*, *CO2_H4M1*, *Q10_CO11*, *Q3_MADS25-2*, *Q3_FT1*, *Q3_CO8*, *CO1*; *green* ), yield-component candidate genes (*brown*); and QTL for spike sterility (qStrlSpk), grain spiculation of inner lateral nerve (qGSILN), spike row number (qRow), are also shown in *dark red*. **Track 2** LocalGEBV estimated using Bayes R, track colour in *blue*. **Track 3** LocalGEBV estimated using rrBLUP, track colour in *green*. **Track 4** GWAS using BLINK, track colour in *orange*. **Track 5** GWAS using FarmCPU, track colour in *purple*. In both **Track 2** and **Track 3**, localGEBV haploblocks were according to LD blocking parameters: LD threshold 𝑟^2^ = 0.1 and marker 𝑡𝑜𝑙 = 3; and shown is the variance localGEBV according to the position of flanking markers within haploblocks. In both **Track 4** and **Track 5**, markers were considered significant according to thresholds after FDR corrections (− log_10_ 𝑃_𝑎𝑑𝑗_ > − log_10_ 0.008 for FarmCPU and − log_10_ 𝑃_𝑎𝑑𝑗_ > − log_10_ 0.004 for BLINK).

Haploblock 2H:b000235 exhibited the highest localGEBV variance (𝑟𝑟𝐵𝐿𝑈𝑃 = 0.0082; 𝐵𝑎𝑦𝑒𝑠𝑅 = 0.0133). This haploblock (spanning 645.10 to 655.33 Mbp) contained all 4 significant markers detected by GWAS, including the most significant marker 2H:AVRIG03047 (− log_10_ 𝑃_𝑎𝑑𝑗_; FarmCPU = 17.6, BLINK = 10.5) at 652.42 Mbp, co-locating with the *VRS1* gene (Komatsuda et al., 2007; Table 1, Figure 4).

Notably, the localGEBV approach identified additional high variance haploblocks significant after Bonferroni or FDR correction that collocated with known genes and candidate genes related to row phenotype. Both rrBLUP and BayesR identified haploblocks 1H:b000049, 3H:b000382, 4H:b000552, and 7H:b001032 co-locating with *VRS3* (associated with grain uniformity; Van Esse et al., 2017; Bull et al., 2017), *HORVU3Hr1G036930* (linked to lateral spikelet fertility; Youssef et al., 2020), and lateral spike fertility (*VRS5* and *HORVU7Hr1G090180*; Ramsay et al., 2011; Youssef et al., 2020) respectively. BayesR uniquely identified haploblock 1H:b000091, co-locating with the QTL associated with “third outer glume” structure (TRD). None of these blocks contained significant markers in the FarmCPU and BLINK analyses.

### Haploblocks demonstrate superior predictability compared to single markers

Using a classical linear model and generalized linear model framework, we aimed to demonstrate that the discrete, categorical haplotype configurations (i.e., discretised localGEBV values) had greater ability to predict row-type phenotype compared to discrete, categorical single marker genotypes. We first conducted principal component analysis on the marker effects from each method of estimating marker effects and conducting GWAS (FarmCPU, BLINK, rrBLUP, and Bayes R). Principal component analysis of marker effect estimates from all methods showed PC1 and PC2 explaining 73.93% of cumulative explained variance, with marker 2H:AVRIG03047 contributing most substantially (17.49%), followed by 4H:AVRIG07945 (5.64%; Figure 5A). Marker 2H:AVRIG03047 was part of the haploblock collocating with the *VRS1* gene and identified as significant by all GWAS and haploblocking strategies (Table 1), whereas marker 4H: AVRIG07945 was only identified in haploblock b000636 using BayesR marker effects.

**Figure 5.**
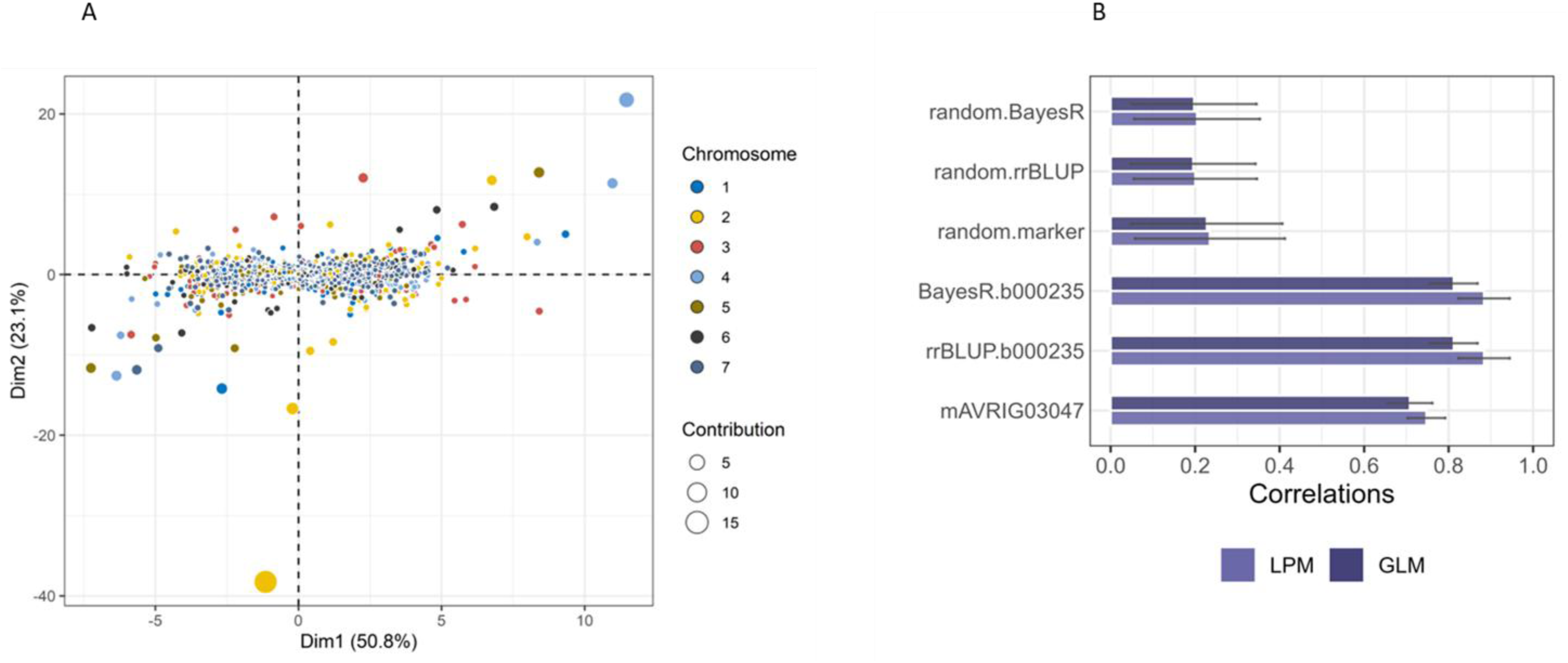
Accuracy of predictors in identifying informative associations with row-spike phenotype. (**A**) Principal component analysis showing the effects contribution of *M* = 9,050 XT SNP marker estimates from FarmCPU, BLINK, rrBLUP, and BayesR. *Colours* represent the 7 barley chromosomes; *size* of each dots represents the SNP effect contribution to the principal components relating to row-spike phenotype. (**B**) Regression analysis of marker genotypes or haploblock haplotype configurations as predictors. Shown is the accuracy calculated as correlations of the predicted value in the testing set; predicted values calculated using ordinary least squares linear probability model (LPM, *light purple*) and logistic regression in the generalised linear model (GLM, *dark purple*). The most significant marker (2H:AVRIG03047) identified by both FarmCPU and BLINK and the haploblock with the highest variance from rrBLUP and BayesR localGEBV estimates (2H:b000235) were analysed in each model and compared with non-significant random marker/haploblock haplotype configurations. Analyses for random marker/haploblock haplotype configurations were repeated 100 times, and the average correlation and standard deviations of the 100 repetitions represented as error bars were shown.

We further evaluated the regression of row-spike status on SNP genotypes or localGEBV haplotype configurations fit as categorical effects for row-spike phenotype prediction accuracy utilising both a linear probability model (LPM) and logistic regression in a generalized linear model (GLM) framework. Configurations were fit as categorical predictors to capture non-additive effects and to avoid utilizing the localGEBV and marker effect values trained on the whole population. This ensured that the predicted effects were specific to the sampled training set. The SNP-only model (mAVRIG03047) used the single most significant marker identified in the FarmCPU/BLINK, while the other two models (rrBLUP.b000235 and BayesR.b000235), incorporate information from haplotypes within the 2H:b000235 (𝑟^2^ = 0.1, 𝑡𝑜𝑙 = 3) simultaneously. The average of the five-fold cross-validation correlations between the predicted phenotype and actual phenotype in the test set were consistently higher for both significant markers and the highest variance haploblock compared to randomly selected markers or haploblocks (Figure 5B). Notably, both rrBLUP and BayesR achieved higher correlation values when using haploblock haplotype configurations as predictors (LPM = 0.88, GLM = 0.81; Root Mean Squared Error; RMSE LPM = 0.07, GLM = 1.78) compared to single marker genotypes (LPM = 0.75, GLM = 0.71; RMSE: LPM = 0.07, GLM = 3.31). The RMSE for the LPM utilising single markers was lower. This is likely a downwards biased estimate because of the violation of ordinary least squares assumptions and should not be utilised to justify a better model fit than the logistic regression model.

### Haplotype analysis reveals distinctive features of localGEBV

We explored the relationship between haploblock variance and the effect of individual haplotypes (Figure 6). In general, as the magnitude of a localGEBV increased, the variance of the corresponding block was greater (Figure 6, Supplementary Figure 3).

**Figure 6.**
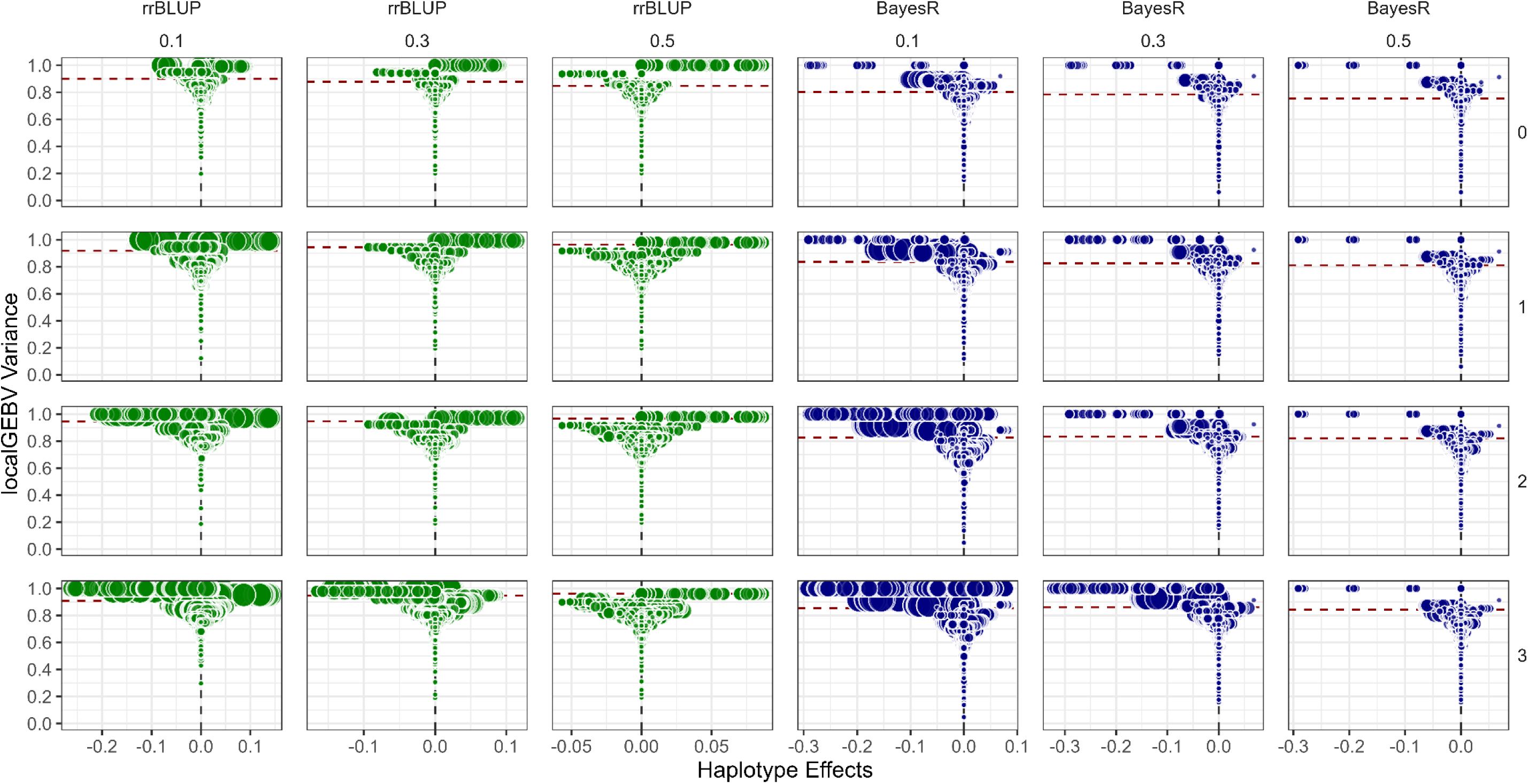
Relationship between haploblock variance and haplotype effects. Haploblock variance was determined from the localGEBV estimates using rrBLUP (*green*) and BayesR (*blue*); plotted as min-max scaled of the *log_10_* transformed variance for visual clarity. In the figure, different LD parameters are shown as facets: *column facet* represented by LD threshold at 𝑟^2^ ∈ {0.1, 0.3, 0.5}; and *row facet* as marker tolerance 𝑡𝑜𝑙 ∈ {0, 1, 2, 3}. *Dashed horizontal lines* represent the average haploblock variance at each LD parameter, where haploblocks above the threshold are considered containing the QTL. *Size* of each circle corresponds to the number of markers within the haploblock.

Detailed haplotype profiling of haploblock 2H:b000235 (*VRS1* region) at higher LD resolution (𝑟² = 0.5, 𝑡𝑜𝑙 = 3) revealed 4 SNP markers spanning 0.88 Mbp (651.54 to 652.42 Mbp) with varying allele frequencies (Figure 7B). From these markers, 30 haplotypes were identified, with 13 occurring at frequencies ≥5 accessions. Two predominant haplotypes emerged as complete allelic reversals: hap1 (TTTT) associated primarily with 2-row phenotype (252) and hap2 (AAAA) with 6-row phenotype (239).

**Figure 7.**
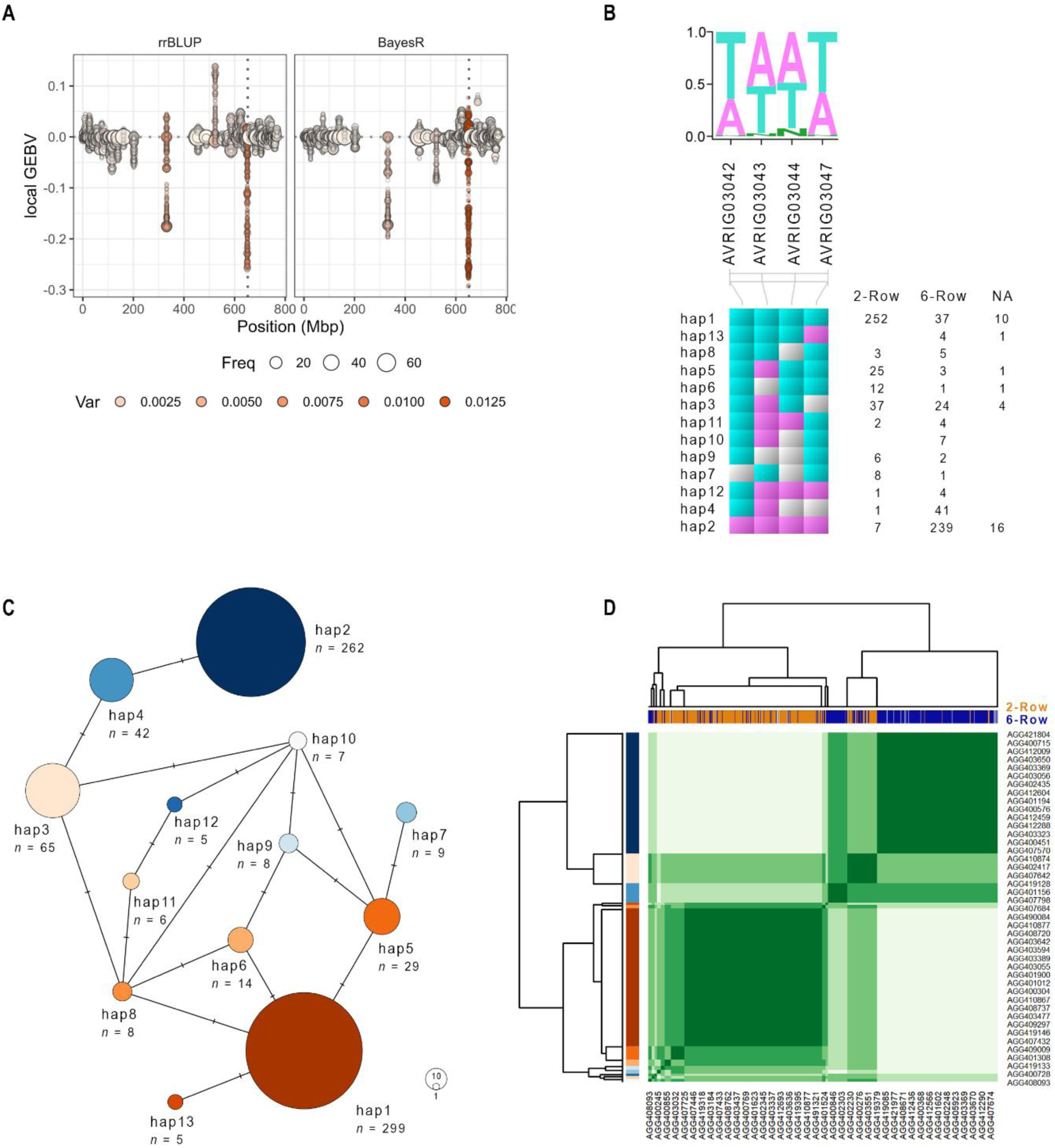
Haplotype analysis. (**A**) Identification of haplotypes from the localGEBV estimates using rrBLUP and BayesR. In the figure, *x-axis* shows the marker physical position (Mbp) of chromosome 2H based on cv Morex assembly v2 from the 40K XT SNP chip; while the *y-axis* corresponds to localGEBV or haplotype effects. The *dotted vertical line* illustrates the position of *VRS1* gene at 652.03 Mbp. Haplotypes associated with row-type phenotypes are determined by the direction of localGEBV estimates, whereby negative localGEBV effects favours 2-row phenotypes while positive localGEBV effects favours 6-row phenotypes. Each *dot* corresponds to a specific haplotype, wherein the *size* of the dots represents the haplotype abundance or the frequency of barley accessions carrying the haplotype; while *orange colour gradient* represents higher haploblock variance. (**B**) Haplotype profile collocating *VRS1* gene. Shown is the 0.88 Mbp finer region within 2H:b00023 defined by four markers (i.e. AVRIG03042, AVRIG03043, AVRIG03044, AVRIG03047; spanning 651.54 to 652.42 Mbp); wherein, arbitrarily, *A* corresponds to the reference allele, *T* corresponds to the alternate allele, and a missing allele is represented by *N*. Marker AVRIG03047 was also the most significant marker identified by FarmCPU (− log_10_ 𝑃_𝑎𝑑𝑗_ = 17.59) and BLINK (− log_10_ 𝑃_𝑎𝑑𝑗_ = 10.49). The *top panel* shows the estimated density of alleles at each four marker positions, while the *bottom panel* shows the 13 haplotypes identified and their frequency of occurrence across barley accessions relative to each row-type phenotype. Only haplotypes with a frequency *of at least* 5 accessions are shown. **(C**) Haplotype network analysis. Shown is a statistical parsimony network (TCS method; Templeton et al., 1992) of the 13 haplotypes. The *size* of the circle is proportional to the number of barley accessions carrying the respective haplotypes, while the *hash marks* indicate the number of mutational steps. *Colour gradient* illustrates the localGEBV or haplotype effects, with *dark orange* and *dark blue* indicating the lowest (i.e. negative) and highest (i.e., positive) haplotype effects, respectively. (**D**) Heatmap of similarity matrix score of the haplotypes among 759 barley accessions. In the figure, barley accessions are separated into two major clusters according to complete linkage hierarchical clustering, wherein the *top colour bar* represents the row-spike phenotypes of the barley accession (i.e., 2-row in *orange* and 6-row in *blue*) while the *left colour bar* represents the 13 haplotypes described in panels B and C.

Haplotype network analysis demonstrated that the direction of haplotype effects aligned with phenotypic expression: negative effects for hap1 is associated with 2-row and positive effects for hap2 is linked with 6-row (Figures 6, 7C); this aligns with the regression analysis in accurately predicting 2-row or 6-row phenotypes utilising the same haplotype effects (Figure 5B). Hierarchical clustering based on similarity matrix scores differentiated distinct clusters corresponding to haplotype-phenotype associations, wherein Australian Grains Genebank (AGG) accessions carrying hap1 had 2-row spikes while AGG accessions with hap2 had 6-row spikes (Figure 7D).

## Discussion

In this study, we demonstrated that localGEBV offers substantial improvements in QTL discovery compared to standard GWAS methods. By integrating LD information with genomic prediction models, the framework effectively partitions the genome into disjointed chromosome segments with estimated haplotype effects (localGEBV) and haploblock variances. This allowed us to leverage local information in the genome enabling more comprehensive detection of genetic loci controlling complex traits. Furthermore, we assessed the impact different marker tolerance and LD threshold blocking parameters had on discovery to provide a general guideline on how to conduct blocking.

### Biological complexity of barley row-type phenotype as a model for QTL discovery

The barley row-type phenotype represents an ideal model for testing QTL discovery methods due to its simple phenotype and semi-complex genetic architecture (Bull, 2015). The morphological distinction between 2-row and 6-row phenotypes is determined by the fertility status of lateral spikelets. In 2-row barley, only central spikelets are fertile and develop grains, while lateral spikelets remain sterile. In contrast, 6-row barley exhibits fertility in both central and lateral spikelets, producing more grains per spike and significantly influencing yield component traits (Youssef et al., 2017) such as thousand kernel weight and kernel number.

The genetic architecture underlying this trait involves a network of co-regulated genes; however, there is still little information on the exact mechanism (Zwirek et al., 2019). While *VRS1* on chromosome 2H serves as the most downstream genetic component (Komatsuda et al., 2007), several other characterised genes regulate lateral spikelet fertility and overall spike morphology, including *VRS2* in 5H (Youssef et al., 2017), *VRS3* (syn. INTERMEDIUM-A) in 1H (Van Esse et al., 2017; Bull et al., 2017), *VRS4* (syn. INTERMEDIUM-E) in 3H (Koppolu et al., 2013; Koppolu et al., 2022), and *VRS5* (syn. INTERMEDIUM-C) in 4H (Ramsay et al., 2011). Contemporary 6-row barley cultivars typically contain natural recessive *vrs1* and *vrs5* alleles (Zwirek et al., 2019).

### Limitations of standard GWAS approaches

Various GWAS methods remain the predominant method for QTL discovery and genetic architecture analysis but face significant challenges including spurious associations, large background noise, and limited power to explain complex trait variants (Wang and Xu, 2019).

Our study utilised a diverse barley panel with pronounced population structure, providing an opportunity to evaluate different mapping approaches with a trait documented to have intricate genetic control.

Despite the significant improvement of statistical methods in both FarmCPU and BLINK that employ multi-locus genome scans (Liu et al., 2016; Huang et al., 2019; Wang and Zhang, 2021), these approaches only detected the major *VRS1* gene, failing to identify other known genes associated with row-type characteristics (Table 1). This limitation exemplifies a common problem in complex trait analysis where primary morphological characteristics are influenced by secondary traits confounded by genetic background (Zwirek et al., 2019) or masked due to epistatic interaction (Ramsay et al., 2011). The low detection power also likely stems from uncompounded effects of co-regulated genetic interactions (Tam et al., 2019), even after controlling for population structure. This can occur when the effect of a QTL is split amongst multiple markers because of extensive LD (correlation) structures between the markers and QTL. Alternatively, each of the multiple markers may only capture a part of the QTL effect if they are only partially correlated with the QTL (Meuwissen and Goddard, 2000; Marchini et al., 2004; Goddard, 2009; Zhang et al., 2010).

Notably, a critical constraint of GWAS is the reliance on stringent significance thresholds, which are implemented to decrease false positives at the expense of decreasing power and increasing the number of false negatives for variants with modest effects. The conventional approach of applying genome-wide significance thresholds (FDR adjustments or Bonferroni-corrected values) prioritizes certainty of associations over discovery breadth. This limits statistical power, excluding true causal variants that fall below the significance threshold (Wang et al., 2016; Tam et al., 2019). A recent alternative to Bonferroni and FDR adjustment on all markers utilized the LD between markers to calculate the true number of effective tests and control type I error (Gao et al., 2008). This would likely increase the power to detect QTL and is based on a similar philosophy to haploblocking, but it would not necessarily recover any split signal like the localGEBV method would. However, we employed a similar approach in a GWAS framework to compute the orthogonal amount of information each block contained to validate the use of haploblock variance as a useful metric in breeding and QTL discovery independent of GWAS. Utilizing this same approach, we were able to increase our power in a GWAS framework as evidenced by the detection of known QTL (Table 1). As one might expect, the relationship between haploblock variance and p-value was quite strong (Supplementary Figure 4), and our approach was able to identify the known QTL despite Bonferroni and FDR p-value correction. Interestingly, the unadjusted p-value distribution was heavily skewed towards 1 (Supplementary Figure 4) with a small increase between 0 and 0.10. The skew towards 1 and small increase near 0 indicated our approach was highly conservative compared to the expected uniform null p-value distribution while also preserving blocks associated with known QTL.

In a trait where multiple genes contribute through intricate regulatory networks, like barley row-type, the statistical stringency employed by classical GWAS methods disproportionately penalize small-effect loci that collectively determine phenotypic expression for complex, polygenic traits. This in turn leads to GWAS having low power to disentangle background noise from true signal (e.g., Figure 4). This phenomenon has been well-documented across multiple crop species. In maize, numerous small-effect loci controlling numerous phenotypic traits were missed by conventional GWAS but enriched with synonymous SNP because of linkage with nearby causal SNP (Wallace et al., 2014). In rice, chlorophyll content and stay-green traits are governed by networks of interacting genes whose individual effects were too small to reach significance thresholds yet collectively explained substantial phenotypic variation (Zhao et al., 2019). Bustos-Korts et al. (2019) demonstrated that barley adaptation traits, such as developmental and row-type traits, are controlled by complex networks of interacting loci whose effects become diluted when analysed individually. However, these effects can be captured by exomic haplotype states due to the larger number of allelic states, supporting our observation that genetic interactions in the *VRS* gene network may go undetected using conventional approaches.

### Methodological advantages of localGEBV

In contrast to standard GWAS, the localGEBV approach demonstrated superior detection power for known genes associated with barley spike row phenotype. This is because the localGEBV haploblocking method prevents overcorrecting p-values as it accounts for non-independency among markers, as previously discussed in GWAS limitations (similar to Gao et al., 2008 methods). The localGEBV method also collates genomic signals that are truly dispersed because of LD in small to moderate effect loci, which maximizes power and simultaneously reduced background noise. This was especially true under the infinitesimal assumptions of rrBLUP, which tended to disperse the signal (e.g., Supplementary Figure 2). Thus, we likely controlled the number of false negatives by collating dispersed QTL signals when accounting for LD structure and preventing the overcorrection of p-values. We demonstrate this by showing that FarmCPU and BLINK detected only the major *VRS1* gene on chromosome 2H, whereas the localGEBV approach additionally identified haploblocks containing other known row-phenotype regulators. These included *VRS3* in 1H (Van Esse et al., 2017; Bull et al., 2017), *VRS5* in 4H (Ramsay et al., 2011), and several annotated QTL (Youssef et al., 2017) that remained undetected by conventional methods (Figure 4).

Notably, the localGEBV methods reduced the background noise compared to the GWAS results from FarmCPU and BLINK, and clearly delineated genomic signals regardless of whether marker effects were estimated using rrBLUP or BayesR. The smoothing of background noise (e.g., Figure 4 comparing variance of localGEBV background to FarmCPU and BLINK background) may be indicative of marker effects from non-QTL regions averaging towards small values or additional shrinkage if the markers were non-informative. The averaging and/or shrinkage would reduce the corresponding haploblock variance when combined into localGEBV utilising LD structure. This demonstrates our method’s ability to leverage localised genomic information to capture QTL signals that would otherwise be lost to background noise and false-positive control. Admittedly, the repressed background noise was initially surprising, because haploblock variance would be expected to scale with the number of markers in the block in a situation where all markers were independent. We demonstrated this to not be the case in Figure 2 comparing number of markers to the haploblock variance and by fitting a generalized additive model. This highlights that using LD to define block structures counteracts this increase and was broadly successful in reducing background noise.

In addition to greater detection power to known QTL, the localGEBV method leverages localised information on LD to fine-tune the region of interest. This relies on historic recombination, drift, and selection patterns that influenced the LD development in the genomic region. In contrast, when interpreting GWAS results, there is no defined region of interest and genome-wide average LD patterns are often utilised to investigate potential QTL or genes of interest. Thus, our methodology helps to accurately assess where a QTL may reside (Figure 3B). However, the choice in haploblock parameters clearly suggested there may be a trade-off in QTL discovery compared to greater genomic resolution to precisely map the QTL. Furthermore, if parameters are too stringent, the QTL may lie outside of the haploblock region even if it is determined to be significant. This is because other markers outside of the block may also be capturing QTL effects even if they are in lower LD with other markers.

A key advantage of the localGEBV approach beyond statistical power and accurately defining QTL regions is its flexibility in accommodating different genomic prediction models to estimate haplotype effects, as exemplified in Figure 4 and 5B. BayesR demonstrated exceptional precision in estimating large-effect QTL, particularly visible in the *VRS1* region. This is not surprising as it aligns with the major role of *VRS1* in the development of the six-rowed spike phenotype controlled by a single recessive allele (*vrs1*), in contrast to the dominant allele for the two-rowed phenotype (Komatsuda et al., 2007). This advantage stems from the model’s ability to accommodate various assumptions in estimating marker effects (Erbe et al., 2012; Moser et al., 2015; Wolc and Dekkers, 2022), leading to improved genome-wide QTL discovery (Kemper et al., 2015; Van Den Berg et al., 2019; Xiang et al., 2021). Conversely, rrBLUP showed advantages in detecting candidate regions with multiple small-effect alleles that cumulatively form a significant signal (Figure 4, Supplementary Figures 2 and 4) and is suggestive of greater applicability for most quantitative traits with near-infinitesimal genetic architecture. However, these candidate regions could not be explicitly verified, because they were not known QTL, and thus more work is needed to determine if rrBLUP truly had greater power than BayesR for small to moderate effect QTL. This model flexibility allows application without prior knowledge of the trait’s genetic architecture, making it suitable for diverse breeding scenarios. For instance, this is evident in the nearby *VRS1* region, where rrBLUP captured higher variance blocks (2H:b000210; Figure 3, Supplementary Figure 2). This region was not found to co-locate with known genes or QTL, suggesting it could represent a potentially new QTL associated with row-type and requires further investigation to determine if it is a new QTL. Thus, our results indicate that the localGEBV method can capture different distributions of marker effects by utilising the infinitesimal assumptions found in rrBLUP.

### LD parameters and genetic implications

The precision of QTL discovery using localGEBV is directly influenced by LD structure within the population. Indirectly, it is also based on marker positions. While we had a high-quality reference map, it may be prudent to ignore marker positions and use the LD patterns (genetic map) to form haploblocks if the reference is poorly constructed. It is also worth noting that we utilized genome-wide blocking parameters, but parameters can also be defined at the chromosomal level.

Our analysis determined that while varying LD parameters affected haploblock size, they did not alter the detection of informative segments for large, known effects (e.g., Figure 3A).

This finding aligns with previous studies concluding that no ‘universal’ or optimal haploblock construction method applied to breeding programs exists (Andrade et al., 2019; Difabachew et al., 2023; Weber et al., 2023). Stringent LD (𝑟^2^ = 0.5) produced smaller haploblocks (often comprised of a single marker) enabling fine mapping of specific genes, while relaxed LD (𝑟^2^ = 0.1) generated larger haploblocks (Figure 3) independent of the genomic prediction method. The ability to choose parameters allows for flexibility and balancing the trade-offs between detection proportion and fine-resolution mapping when conducting QTL discovery studies. This is a clear advantage not currently found in most, if not all, GWAS methods without utilising different priors on the underlying genetic architecture. In conjunction with likely lower power to detect low to moderate effect QTL based on our results, the area to search for QTL is often defined by genome-wide average LD patterns. The localGEBV method overcomes both challenges.

The biological significance of haploblock size warrants consideration. Smaller haploblocks may signify high LD between markers and the QTL (Andrade et al., 2019), which facilitates precise candidate gene identification for post-GWAS analysis (Clark, 2004). On the other hand, larger haploblocks likely capture important biological mechanisms including local epistasis, pleiotropy (Kouyos et al., 2007; Zhang et al., 2014; Jiang et al., 2018), and adaptive mechanisms from balancing selection, mutation, and genetic drift (Barton, 2010). We did not observe this in our case, but we utilised a genetically intricate trait rather than fully Mendelian with substantial non-additive effects between a low number of loci.

Additionally, information on recombination hotspots at haploblock boundaries (Kim et al., 2018; Epstein et al., 2023) can be valuable for breaking unfavourable linkage drag in breeding programs (Bohra et al., 2022) or stacking favourable haplotypes. Utilizing haploblocks provides an interesting advantage when introgressing new QTL (haplotypes) for a complex population structure. Utilising localGEBV from haploblocks provides a “genetic insurance policy” by co-selecting target genes with their local genetic background, thereby ensuring more predictable and stable trait expression when introgressed into new cohorts. The latter point is especially important in genetic improvement to maintain diversity and construct the “ultimate genotype” (Cole and VanRaden, 2011; Kemper et al., 2012; Hayes et al., 2024). The latter suggests our results likely have implications in breeding programs, crossing designs, and genomic improvement. This is comparable to GWAS and genomic selection sharing the same underlying framework and considerations, so it is natural to extend our framework and its implications to genomic improvement. In such a case, selection of blocks based on the haploblock variance or utilising the number of blocks capturing a set proportion of the genetic variation could be utilised in breeding applications.

Lastly, it is important to note that constructing haploblocks based on LD is not always possible. When constrained by LD structures from diverse populations (Abdellaoui et al., 2023) like those commonly found in diversity panels, when SNP density is not homogeneous (Li et al., 2021), or when markers are extremely dense with common alleles mixed with rarer alleles (Taliun et al., 2014), alternative haploblock construction methods such as sliding window approaches (Weber et al., 2023), gene boundary-based collapsing methods (Dering et al., 2011), or incremental haploblock partitioning (Taliun et al., 2014) could be more advantageous. However, the implications of these methods to QTL detection in the localGEBV framework has not yet been explored.

### Biological insights from localGEBV features

The localGEBV approach provides deeper understanding of trait genetic architecture through two key features:

### Haploblock variance

Haploblocks with high variances (greater than the mean haploblock variance) were consistently associated with substantial haplotype effects, representing true biological signals rather than artifacts of segment size because size was not correlated with variance (Figure 2 and 6). This variance metric offers advantages over conventional p-values in GWAS, which provides statistical significance but often lack informative biological meaning (Visscher et al., 2017), may suffer from false associations (Yang et al., 2010; Wray et al., 2013) and do not provide a definitive region to explore QTL. In addition, GWAS may also only capture part of the QTL signal or share its effect with nearby markers in high LD with the QTL (Meuwissen and Goddard, 2000; Marchini et al., 2004; Goddard, 2009; Zhang et al., 2010); however, this is dependent on the prior assumptions of marker effects, towards which our approach is more robust. By shifting focus from marker significance thresholds to haploblock variance or testing haploblock variance for significance, the localGEBV approach diminishes the problems associated with multiple testing that also makes small-effect loci effectively undetectable. The reduction in stringency is because there are often far less haploblocks than markers, so less stringency on type I error control is needed. The reduction in the number of tests is appropriate, because markers are not independent and Bonferroni or FDR corrections assume test independence (Gao et al., 2008). Variance patterns can indicate a high degree of polymorphism in regions under selection pressure or fixation (Maynard and Haigh, 2007), absence of selective interactions (Otto and Barton, 1997), or enriched mutations affecting trait heritability (Sullivan et al., 2023).

### Haplotype effects

Haplotypes provide superior predictive ability compared to single markers. Our regression analyses demonstrated that the haplotype configurations within a particular, high-variance haploblock serve as better predictors of phenotypic traits than individual SNP genotype configurations. This highlights the importance of considering cumulative genetic effects in tightly linked regions (Meuwissen and Goddard, 2000), rather than isolated markers as single markers can only capture a limited amount of genetic variation.

The choice of estimating SNP effects (Step 2) to calculate localGEBV provides flexibility of this approach. While BayesR was able to confirm known large effect *VRS1* region (Figure 3 and 7A; Supplementary Figure 2), complex traits will not have this type of architecture and rrBLUP or other similar infinitesimal models is a good default model.

Lower-frequency alleles with considerable effect can be successfully captured as a single unit within the haploblock (Berdan et al., 2023), enabling exploration of local allelic interactions and epistasis (Charlesworth, 1974). The localGEBV method may help to capture these rare effects, but only if there are QTL and markers that have similar frequencies because LD is dependent on similar allele frequencies.

Lastly, haplotype effects also signify directionality (Han and Pan, 2010) in relation to a reference haplotype, similar to the allele substitution effect of the alternative allele relative to the reference allele. Our analysis confirmed that negative-effect haplotypes were associated with 2-row phenotypes and positive-effect haplotypes with 6-row spikes (Figure 7A and 7D).

### Limitations of localGEBV

The localGEBV method represents a promising leap forward from a single marker to LD-based haplotype approach. However, the implementation we present here is based solely on additive genetic effects and aggregating dispersed QTL signals using LD. Regarding the latter, this means that haploblocks are based on historic selection, drift, and recombination patterns that influence the development and decay of LD (Slatkin, 2008). However, if intense selection and drift were driving quick formation of LD, it could indicate that haploblock boundaries are not reflective of future recombination “hotspots.” In such a case, the localGEBV are not representative of regions that will be inherited more often than expected, but may still aggregate QTL effects dispersed across multiple, highly correlated markers (i.e., statistical advantages rather than biological interpretation). Thus, biological interpretation of haploblock boundaries is determined on the specific population that is analysed, its population structure, and recent selection history.

Furthermore, while only additive genetic effects are considered here, non-additive genetic effects, such as dominance and epistasis, could be incorporated into the calculations to compute local genomic estimated genotypic values (localGEGV) rather than localGEBV. It is conceivable that dominance effects could be “solved” for individual loci much in the same way additive marker effects are derived from Genomic Best Linear Unbiased Prediction using dominance or genotypic value relationship matrices (e.g., Vitezica et al., 2017; Liu et al., 2022). The local genomic estimated dominance deviation values for a haploblock would then simply be the sum of the estimated dominance deviation values across the loci in the block and the localGEGV the sum of local genomic estimated dominance deviation values and localGEBV. Bayesian methodologies to directly compute dominance deviation and genotypic values have similar interpretations to directly computing additive marker effects using Bayesian alphabet methods (e.g., Wellmann and Bennewitz, 2012). Vitezica et al. (2017) have similarly proposed methods to compute orthogonal epistasis effects, which represents an opportunity to exploit epistatic interactions. However, how epistatic interaction effects would be incorporated from epistatic relationship structures, or even a direct interaction model (Bayesian or otherwise) remains an active area of exploration.

### Implications for crop breeding

The localGEBV approach offers significant advantages in QTL discovery that could be used in genetic improvement strategies in crops. By exploiting LD blocks, breeders can determine both global and local genetic correlations between multiple traits (Visscher et al., 2017; Olasege et al., 2022), identify sources of genetic control under pleiotropic influence (Watanabe et al., 2019), and select favourable haplotype configurations. The approach effectively identifies compound-effect loci (Bevan et al., 2017) that influences complex traits. The approach is particularly valuable for traits that are difficult to phenotype; for example, fertility in barley is difficult to phenotype because fertility of lateral spikelets drives row-type morphology. Consequently, fertility can influence other yield components such as grain number and thousand kernel weight.

As an example of a useful application, haploblocks that may benefit two unfavourably correlated traits can be identified to guide parental selection and stack haplotypes that may increase the rate of long-term genetic gain compared to utilising genome-wide GEBV. In essence, precise parental combinations and mating pathways can be identified to ensure long-term, consistent genetic gain in unfavourably correlated traits. This would be at odds with multi-trait selection in traditional genomic prediction frameworks, because multi-trait selection is expected to exhaust favourable genetic covariation between the traits, leaving only neutral or unfavourable genetic covariation (Itoh, 1991). The latter would make genetic progress for both traits increasingly more difficult over time (Itoh, 1991).

In practice, linkage drag between haplotypes pose a challenging task in breeding. Haplotype boundaries signifying recombination hotspots can be broken down using targeted recombination approach. On the other hand, for a more deployable application, localGEBV information can be used to develop haplotype markers, especially for qualitative or oligogenic traits such as disease resistance.

The ability to map haploblocks and identify specific haplotypes associated with traits provides a powerful framework for unravelling genetic architecture that can be used to advance crop breeding efforts. While we used a diverse germplasm with unique population structure in this paper, the localGEBV method can also be applied in commercial breeding populations or other experimental populations such as MAGIC or NAM, showing higher detection ability (e.g., Varshney et al., 2021; Brunner et al., 2024; Tong et al., 2024; Akinlade et al., 2025; Alahmad et al., 2025; Roy et al., 2025; Vo Van-Zivkovic et al., 2025; Aldiss et al., 2025). These insights are essential for developing haplotype-based selection strategies that assemble beneficial allele combinations and optimise genotype configurations (Hayes et al., 2024), which would complement gene networks and genotype-to-phenotype systems approaches (Cooper et al., 2009; Kemper et al., 2012; Villiers et al., 2024), and improve genomic predictions (Schopp et al., 2017) in crop breeding programs.

To summarize the benefits to QTL discovery and breeding programs, the haploblock approach represents an innovative compromise between GWAS and genomic selection, combining the strengths of both methodologies while addressing their respective limitations. While GWAS provides precise identification of QTL underlying traits of interest, its applications remain largely confined to pre-breeding activities with limited direct utility for practical breeding programs. Conversely, genomic selection effectively estimates genome-wide breeding values for direct use by breeders but offers limited insights into the underlying genetic architecture of target traits and can severely impact diversity with intense selection. The localGEBV method bridges this gap by simultaneously enabling the dissection of genetic control mechanisms and the identification of genomically-informed selection units (haploblocks) directly applicable in breeding programs. The flexibility to adjust block size according to specific breeding objectives to reflect LD structures further enhances the practical utility of this approach, allowing breeders to optimize the balance between genetic resolution and selection efficiency. Such dual functionality positions haploblocks as a powerful tool for translating genomic discoveries into actionable breeding strategies.

Future research should explore extended genomic prediction models to capture rare variants, non-additive or epistatic interactions, leverage advancements in SNP arrays or whole genome sequencing to improve the haplotype blocking algorithm, haplotype phasing accurately, and explore the applications of localGEBV in selection under extensive genotype-by-environment interactions and multi-trait selection.

In conclusion, we demonstrated the localGEBV methodology is a powerful tool for QTL discovery. The localGEBV method was able to detect more known QTL with clearer signals compared to other GWAS methods. Furthermore, it alleviated or eliminated many of the limitations associated with multiple-testing, background noise, dependence on prior marker distribution (genetic architecture) assumptions, and defining regions of interest for candidate gene/QTL discovery based on significant associations. The localGEBV method was able to locate the largest effect QTL of interest with better prediction accuracy in a genomic selection context and was able to identify other known QTL as well as several putative QTL that FarmCPU and BLINK could not identify. When utilised in a genomic prediction or QTL context, the localGEBV method was robust to *a priori* assumptions of the genetic architecture, with near equivalent phenotypic prediction accuracy from a large effect QTL using an infinitesimal modelling approach or marker prior mixture approach to marker effects.

## Methods

### Barley phenotypic and genotypic datasets

We analysed *N* = 812 barley accessions with complete genotypic and spike row morphology data acquired from the AGG, representing global diversity originating from 80 countries. Accessions were genotyped using the Illumina Infinium wheat-barley 40K XT SNP chip array (Keeble-Gagnère et al., 2021), yielding *M* = 12,539 SNP markers. Genotype calls were in 0/1/2 format (homozygous reference, heterozygous, homozygous alternate) relative to the cv Morex assembly IBSC v2.0.

Data filtering excluded unmapped markers and applied the following criteria using SelectionTools (Hofheinz and Frisch, 2014): >20% missing data, <1% expected heterozygosity, and <1% minor allele frequency. This resulted in *M* = 9,050 mapped markers for analysis. Accessions with >20% missing marker data were also excluded, leading to *N* = 790 accessions remaining.

Spike row descriptions obtained from AGG classified accessions as 2-rowed (373) or 6-rowed (384). For simplicity, we excluded 33 accessions that were described as ‘irregular’ row-type (Åberg and Wiebe, 1945), had missing information, or had ‘mixed’ designations. After exclusion of accessions with ambiguous designations, the dataset with row-type for QTL discovery analyses comprised *N* = 759 barley accessions. All analyses were performed using R version 3.6.3.

### Population genetic structure and linkage disequilibrium analysis

We performed classical multidimensional scaling analysis of Roger’s genetic distance using SelectionTools on the *M* = 9,050 mapped markers to infer population structure across the *N* = 790 barley accessions prior to filtering accessions for ambiguous row-type phenotypes. A *k-means* clustering algorithm categorised accessions into clusters, with the optimal number (*clusters* = 3) determined according to the majority of the criterion indices from the NbClust package (Charrad et al., 2014).

For linkage disequilibrium (LD) assessment, we calculated the squared pairwise correlations (𝑟^2^) between intrachromosomal SNP pairs. The LD decay was evaluated by analysing average 𝑟^2^ between adjacent SNP as a function of physical distance, with positions aligned to the reference cv Morex assembly IBSC v2.0 as in the 40K XT SNP chip array.

### QTL discovery approach

#### ‘Local’ GEBV approach

The localGEBV approach consists of three key steps:

*Step 1: Haploblock construction.* Genome-wide markers were first organised into haploblocks using an LD-based approach (Supplementary Figure 5), which provides insight into physical recombination events. Due to its reduced dependence on allele frequencies and independence from finite population size, we used the squared correlation coefficient, 𝑟^2^, as the LD measure following methodologies described by Voss-Fels et al. (2019).

Given the LD characteristics of our barley population, we tested multiple blocking criteria: 𝑟^2^ ∈ {0.1, 0.3, 0.5} and marker tolerances, 𝑡𝑜𝑙 ∈ {0, 1, 2, 3}, resulting in 12 LD blocking parameter combinations. The algorithm scanned each barley chromosome independently to group SNP markers into blocks based on the LD threshold and tolerance parameters. To make blocks, the 𝑟^2^ measurement between the last marker in a block (or the first marker in a new block) and the adjacent marker was evaluated against the specified threshold to determine whether the marker would be included in the haploblock. If the 𝑟^2^ value was below the threshold, the block would either be terminated, or the tolerance counter would be incremented by one. The marker tolerance parameter specified how many markers could fail the threshold check before a block was terminated. Specifying a tolerance greater than zero allows the algorithm to handle issues in the data sequencing, allele calls, or issues with the reference genome construction or reference alignment. A tolerance of 0 build would build haploblocks strictly based on pair-wise marker LD. A block may be composed of a single marker if it is not in LD with other markers, but is notably still considered a block. The algorithm continued until all markers were grouped into blocks.

*Step 2. Estimation of SNP effects.* We tested two models to estimate SNP effects for comparison: rrBLUP and BayesR. Following the approach described by Voss-Fels et al. (2019), the rrBLUP method was implemented in SelectionTools using:

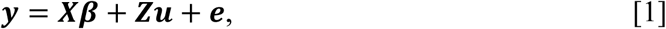

where 𝒚 is the observation vector of row-type (2 or 6) phenotypic values for *N* genotypes, 𝑿 is the *N* × *p* design matrix allocating *N* phenotypes to *p* fixed effects, 𝜷 is the size *p* vector of fixed effects, 𝒁 is the *N* × *M* design matrix allocating phenotypes to the random effects of *M* SNP markers, 𝒖 is the vector of *M* SNP effects, and 𝒆 is the random residual errors. It is assumed that 𝒖 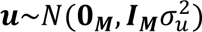 and 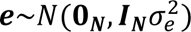; where 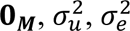 and ***I*** are a vector of zeros of length *M*, the additive genetic marker variance under infinitesimal assumptions, the residual variance, and identity matrices of size 𝑀 and 𝑁, respectively. The rrBLUP solution for the *M* SNP marker effects, *û* (Supplementary Figure 5), can be solved as:

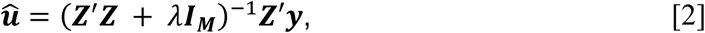

where 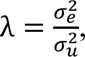 estimated in the rrBLUP software package, represents the ratio between the residual and SNP marker variances.

BayesR (Erbe et al., 2012; Moser et al., 2015) was implemented using GCTB v2.0 (Genome-wide Complex Trait Bayesian analysis, https://cnsgenomics.com/software/gctb) following the approach described by Kemper et al. (2015). The model was fit following the model in Equation 1 and used the same 𝒁 design matrix. However, BayesR assumes *a priori* that SNP effects, 𝒖, are drawn from a mixture of *K* normal distributions with mean 0 and variances 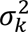, with mixture probabilities defined by vector 𝜸 = [𝛾_1_, …, 𝛾_𝐾_]. At each iteration of the Gibbs sampler, the effect of SNP *m* is updated by first computing its posterior probabilities of membership in each mixture, conditional on the current effects of all other SNP and mixture probabilities in 𝜸. A component assignment 𝑏_𝑚_ is then sampled from the posterior probabilities of membership to each mixture, and the SNP estimate, *û*_𝑚_, is drawn from the corresponding normal distribution. After all SNP have been updated, the mixture proportions, 𝜸, are resampled from a Dirichlet distribution, with 𝜸∼Dirichlet(𝛂 + 𝜹), where 𝛂 is the vector of prior counts and 𝜹 are the counts of SNP assigned to each of the *K* mixtures in an iteration of the Gibb’s sampler.

In our implementation, we set 𝐾 = 4 mixture components corresponding to zero-mean normal distributions with variance 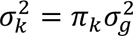 and 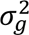 is the total additive genetic variance. The prior mixture probabilities were defined 𝜸 = [0.95, 0.03, 0.01, 0.01] and the prior counts were defined 𝜶 = 𝟏_𝑲_, a vector of ones of length *K*. In other words, while the prior on an individual SNP effect distribution was informative, the prior on the overall genome-wide allocation to each mixture was left uninformative and the posterior was driven completely by the data. We ran 25,000 iterations of the Gibbs chain after 5,000 burn-in iterations.

*Step 3. Estimation of localGEBV and variance.* Linear contrasts of the estimated SNP effects were then specified to aggregate estimated SNP effects within each haploblock, yielding the estimated localGEBV of each individual at each haploblock (Kemper et al., 2015) as follows:

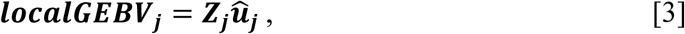

Where 𝒁_𝒋_ = 𝒁_𝒎𝒓:𝒎𝑹_ is the design matrix of the marker genotypes (0/1/2 format for all *N* individuals) comprising haploblock *j* of *J* total haploblocks and *û*_𝒋_ = *û*_𝒎𝒓:𝒎𝑹_ denotes the vector of estimated SNP effects spanning from the first marker (𝑚_𝑟_) to the last marker (𝑚_𝑅_) in haploblock *j* (Supplementary Figure 5). The 𝒍𝒐𝒄𝒂𝒍𝑮𝑬𝑩𝑽_𝒋_is the vector of 𝑁 localGEBV values at block 𝑗, similar to the vector of GEBV from a standard genomic selection model utilizing the whole genome. Estimated SNP effects were obtained from Equation 2 in the rrBLUP implementation or from the posteriors of the BayesR implementation of Equation 1 to compare the two methods.

Subsequently, the variance of the estimated localGEBV effects (haploblock variance) across individuals in haploblock *j* was defined to be:

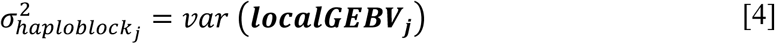

Where 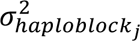 is the variance of the localGEBV at haploblock *j* and 𝒍𝒐𝒄𝒂𝒍𝑮𝑬𝑩𝑽_𝒋_ is the vector of *N* estimated localGEBV effects for all *N* individuals at block *j* (Supplementary Figure 5). The variance represents the population variance calculation on the vector of observed localGEBV values. In standard genomic prediction, 𝑣𝑎𝑟(𝒍𝒐𝒄𝒂𝒍𝑮𝑬𝑩𝑽_𝒋_) is analogous to 𝑣𝑎𝑟(𝑮𝑬𝑩𝑽) in the decomposition, 𝑣𝑎𝑟(𝑩𝑽) = 𝑣𝑎𝑟(𝑮𝑬𝑩𝑽) + 𝑃𝐸𝑉(𝑮𝑬𝑩𝑽), where 𝑣𝑎𝑟(𝑩𝑽) is the variance of true BV, (i.e., the additive genetic variance in a genomic BLUP, GBLUP, 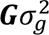), 𝑣𝑎𝑟(𝑮𝑬𝑩𝑽) is the variance of the estimated genomic BV captured by the data, and 𝑃𝐸𝑉(𝑮𝑬𝑩𝑽) is the prediction error variance of the estimated BV. Thus, the haploblock variance represents the additive genomic variance attributable to a genomic segment that can be captured by the data, rather than the total underlying genetic variance of the segment (Mrode and Pocrnic, 2023). Here, we use the haploblock variance as a metric for prioritising chromosome segments containing QTL, conceptually analogous to how locus-specific additive variance is used to prioritise important loci in GWAS. Haploblocks have previously been considered QTL if the variance surpasses the overall mean haploblock variance (Kemper et al., 2015; Yadav et al., 2025).

To assess whether high variance haploblocks were significantly different from the null variance assumption, we calculated probability values (p-values) using a chi-squared distribution with 1 degree of freedom. The test statistic for a given block was as follows:

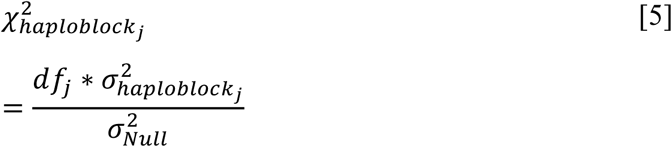

where 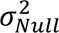 was calculated by dividing the estimated additive genetic variance from GBLUP by the number of blocks (𝐽 = 1096), indicating an infinitesimal assumption where additive genetic variance is split equally among blocks. The 𝑑𝑓 was calculated as the effective number of markers in a block (i.e., the independent and orthogonal pieces of information) by using eigen decomposition on the genotype correlation matrix of markers corresponding to block 𝑗. The number of eigenvalues that explained at least 90% of the variation was considered the effective number of markers in the block, similar to Gao et al. (2008). The resulting p-values were Bonferroni and FDR-corrected and reported. The p-values were only generated for the set of LD parameters (𝑟^2^ = 0.1 and 𝑡𝑜𝑙 = 3) that maximized haploblock variance at the known *VRS1* locus.

To explore the relationship of haploblock variance with size and haplotype effects, a generalized additive linear regression model, regressing haploblock variance on the number of markers in the haploblock or the haplotype effect was performed using the R package mgcv (Wood, 2001) to capture higher degree polynomial trends in the block variance and block size relationship. Each of the 24 sets of LD blocks constructed from the combinations of 𝑟^2^ and tol parameters were analysed independently.

#### Comparative genome-wide association studies and co-location analysis

For comparative GWAS, we used the same genotype matrix in Equation 1 in two models implemented in Genome Association and Prediction Integrated Tool (GAPIT) v3 in R (Wang and Zhang, 2021): Fixed and random model Circulating Probability Unification (FarmCPU; Liu et al., 2016) and Bayesian-information and Linkage-disequilibrium Iteratively Nested Keyway (BLINK; Huang et al., 2019). FarmCPU iteratively incorporates fixed and random effects, whereas BLINK uses only fixed effects. Unlike the standard single-locus mixed linear model, which tests markers one at a time in a one-dimensional genome scan, both FarmCPU and BLINK perform multi-dimensional genome scans that estimate multiple marker effects simultaneously while controlling for population structure.

To account for population stratification, we fitted 15 principal components (explaining ∼80% cumulative variance) as covariates. To control for false discovery rate, significant markers were identified following the built-in *P* value thresholds (− log_10_ 𝑃_𝑎𝑑𝑗_ > − log_10_ 0.008) for FarmCPU and − log_10_ 𝑃_𝑎𝑑𝑗_ > − log_10_ 0.004) for BLINK).

Significant SNP markers identified by FarmCPU and BLINK were then compared with haploblocks showing significant haploblock variance from rrBLUP and BayesR with co-location. Co-location was based on the physical position according to the reference cv Morex assembly IBSC v2.0 described in the 40K SNP chip array. We cross-referenced results with known genes and QTL associated with row phenotypes, inflorescence development, fertility of spikelets, overall spike morphology, and agronomic and yield component traits (Komatsuda et al., 2007; Ramsay et al., 2011; Koppolu et al., 2013; Bull, 2015; Youssef et al., 2017; Van Esse et al., 2017; Bull et al., 2017; Koppolu et al., 2022).

### Assessment of detecting true haploblocks

First, to summarise the estimates of SNP effects obtained from FarmCPU, BLINK, rrBLUP, and BayesR, we calculated their effect contributions in the principal components using the factoextra (Kassambara and Mundt, 2016) R package.

To evaluate the accuracy of detecting a true QTL or haploblocks, we cross-validated the population using an 80:20 split (training set of 589; testing set of 148) in a 5-fold scheme. Detection accuracy, 𝑟, defined as the average from the 5-fold correlations between the observed phenotypic values (*y*) and predicted values (*ŷ*) in the testing set. Specifically, we tested the predicted effects of the haploblock with the highest variance identified by rrBLUP and BayesR to predict the row-type phenotype compared to the most significant markers identified by both FarmCPU and BLINK. This was done by fitting the discrete, categorical haplotype configurations and discrete, categorical marker genotype configurations as fixed effects in a LPM and in a logistic regression within a GLM framework in R. In the LPM, estimation was performed by ordinary least squares, assuming the residual was normally, independently, and identically distributed as an approximation because of the binary nature of the row-type trait. In the logistic regression in the GLM framework, the same linear predictor from the LPM was linked to the binomial distribution of the row-type trait via the logit link function, thereby mapping the predictor to the probability scale.

For comparison, we repeated the analysis with random markers (i.e., non-significant) and random haploblocks (i.e., with localGEBV variance below the threshold). Random selection was repeated 100 times, and the mean correlation, 𝑟, and standard deviations (standard error bars) of the 100 iterations were reported.

### Haplotype analysis

To explore the two unique features of localGEBV (i.e., haploblock variance and haplotype effects), we examined the specific haplotypes associated with the trait. As an illustrative example, the haploblock with the highest variance was first subjected to statistical parsimony haplotype network analysis (TCS method) implemented in POPART (Leigh et al., 2015). Each haplotype, defined by a series of SNP alleles within the haploblock, was represented as a node in the haplotype network. Using these haplotypes, we calculated pairwise similarity scores based on the series of SNP alleles or haplotypes between accessions using FlapJack (Milne et al., 2010) and subsequently performed complete linkage hierarchical clustering of barley accessions. This enabled us to assess the relationship between haplotype similarity and row-spike phenotypes.

## Acknowledgements

We acknowledge Dr Sally Norton and the team at the Australian Grains Genebank for facilitating the acquisition of barley germplasm collection associated accession information; Dr Kerrie Forrest and the team at Agriculture Victoria for genotyping using the 40K Infinium XT SNP Chip; Dr Michael Groszmann at GRDC for discussions on the practical applications of localGEBV method. ED was supported by the Queensland Government’s Industry Research Fellowships Program. LTH was supported through an ARC Future Fellowship (FT220100350). WS and VP were supported as Adjunct Postdocs of the Australian Research Council Training Centre in Predictive Breeding for Agricultural Futures (IC230100016).

## Author contributions

Conceptualization, E.D., K.V., L.H., and B.H.; data analysis and writing – review and editing, W.S., E.D., V.P., and S.Y.; Funding acquisition, E.D. All authors have read and approved the final manuscript.

## Pre-Print Server

The article was submitted to the pre-print server, bioRxiv DOI: https://doi.org/10.1101/2025.08.28.672830

## Supplementary Files

**Supplementary Figure 1.**
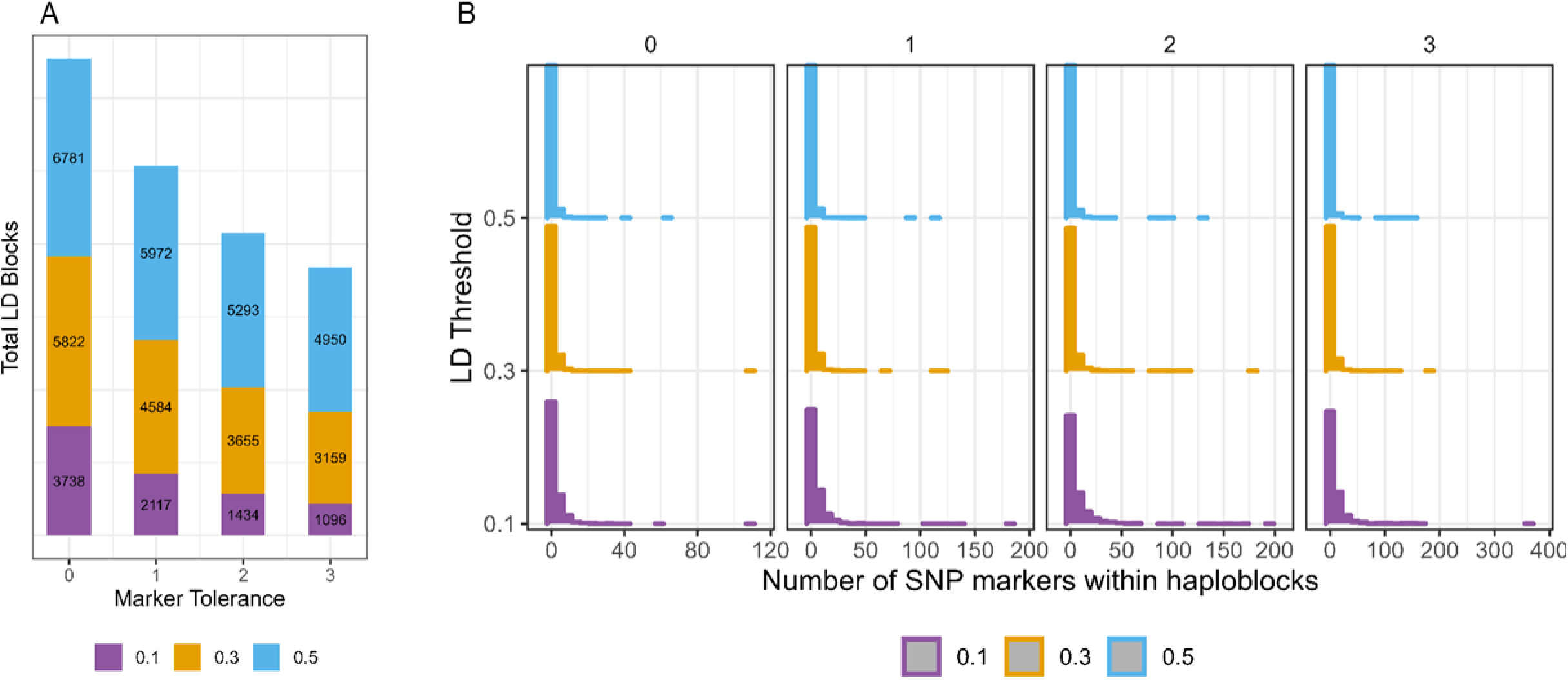
LD block construction and its relationship with block size. SNP markers are grouped according to intrachromosomal pairwise LD based on the different LD-blocking parameters (LD threshold 𝑟^2^ ∈ {0.1,0.3,0.5}; and marker 𝑡𝑜𝑙 ∈ {0,1,2,3}). (**A**) The number of total blocks for each combination of marker tolerance (x-axis) and LD threshold (colours and y-axis) are presented. (**B**) The distribution of the number of markers in each block for each combination of marker tolerance (x-axis) and LD threshold (colours and y-axis) are presented.

**Supplementary Figure 2.**
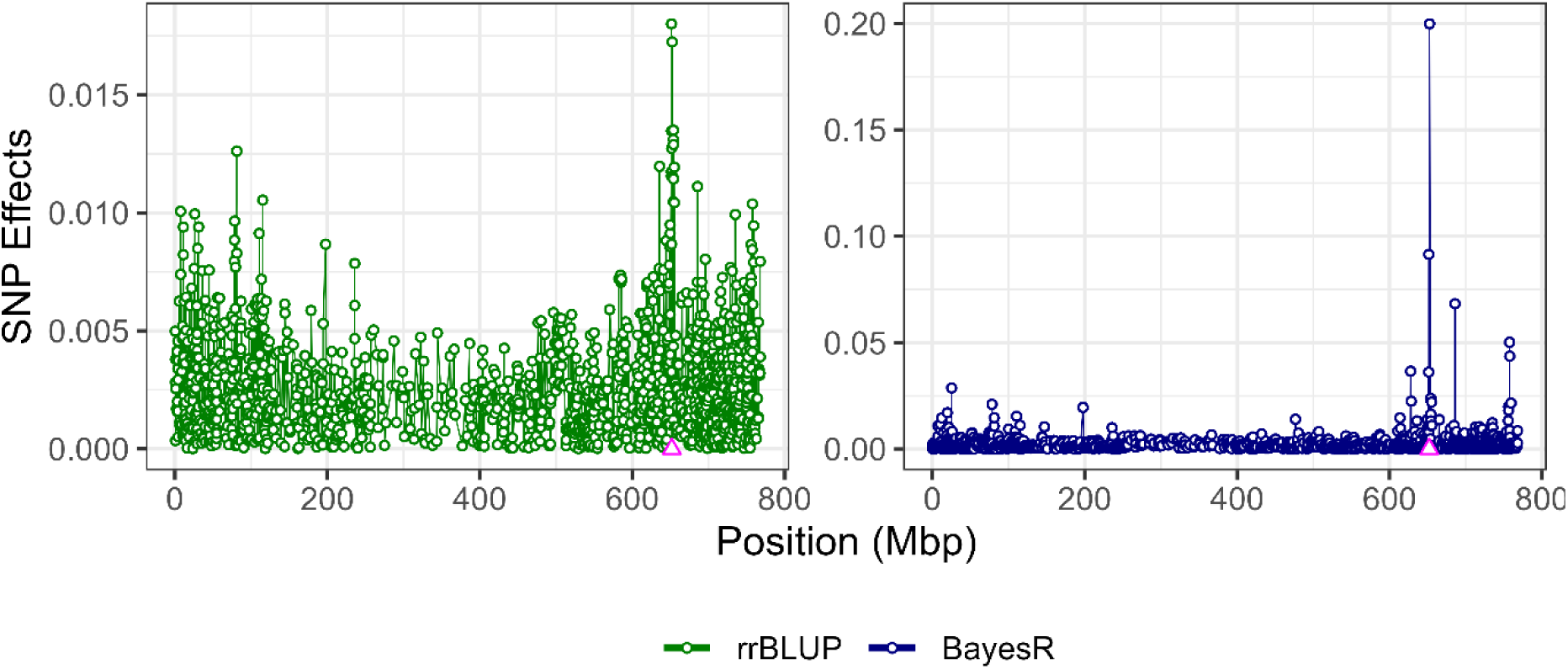
SNP effects estimated by rrBLUP and BayesR for row-type spike architecture in chromosome 2H. Shown is the absolute value of SNP effect estimates, note the change of *y-axis* scale for rrBLUP and BayesR. The *VRS1* gene (marked as Δ) is located at 652.03 Mbp based on the XT SNP chip physical position using cv Morex assembly v2

**Supplementary Figure 3.**
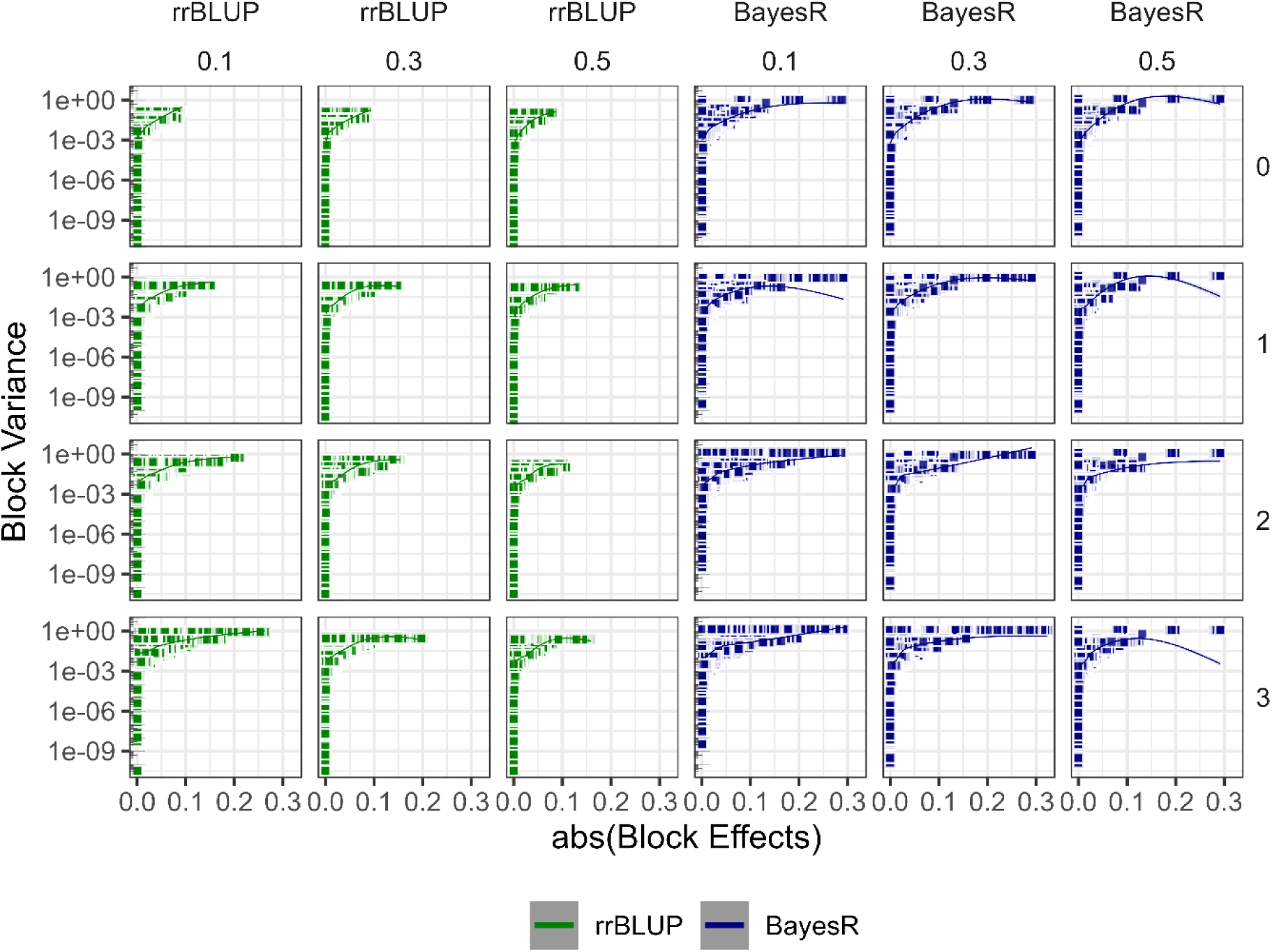
Relationship between haploblock variance and haplotype effects using generalised additive model. Haploblock variance was determined from the localGEBV estimates using rrBLUP (*green*) and BayesR (*blue*); and plotted as min-max scaled of the log_10_ transformed variance for visual clarity. Haplotype effect is equivalent to localGEBV, which is the sum of SNP marker effects within haploblocks. In the figure, different LD parameters are shown as facets: *column facet* represented by LD threshold at 𝑟^2^ ∈ {0.1,0.3,0.5}; and *row facet* as marker 𝑡𝑜𝑙 ∈ {0,1,2,3}.

**Supplementary Figure 4.**
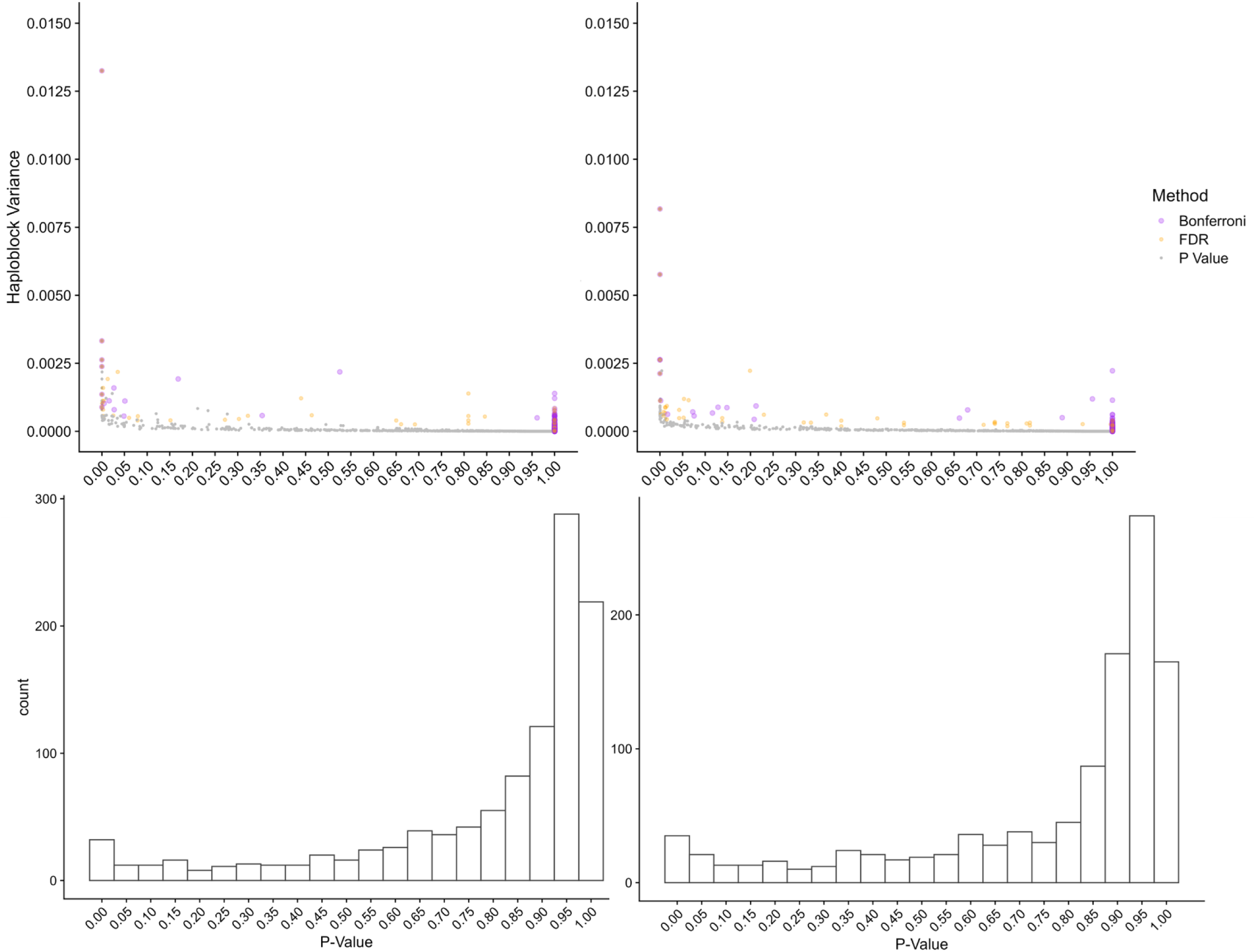
**Characterisation of the distribution of p-values and their relationship to haploblock variance**. (**A**) The relationship between haploblock variance calculated using BayesR, p-values, and p-values adjusted with Bonferroni or False Discovery Rate (FDR) multiple testing correction. (**B**) The relationship between haploblock variance calculated using rrBLUP, p-values, and p-values adjusted with Bonferroni or FDR multiple testing correction. (**C**) The distribution of unadjusted p-values calculating haploblock variance with BayesR marker effects. (**D**) The distribution of unadjusted p-values calculating haploblock variance with rrBLUP marker effects.

**Supplementary Figure 5.**
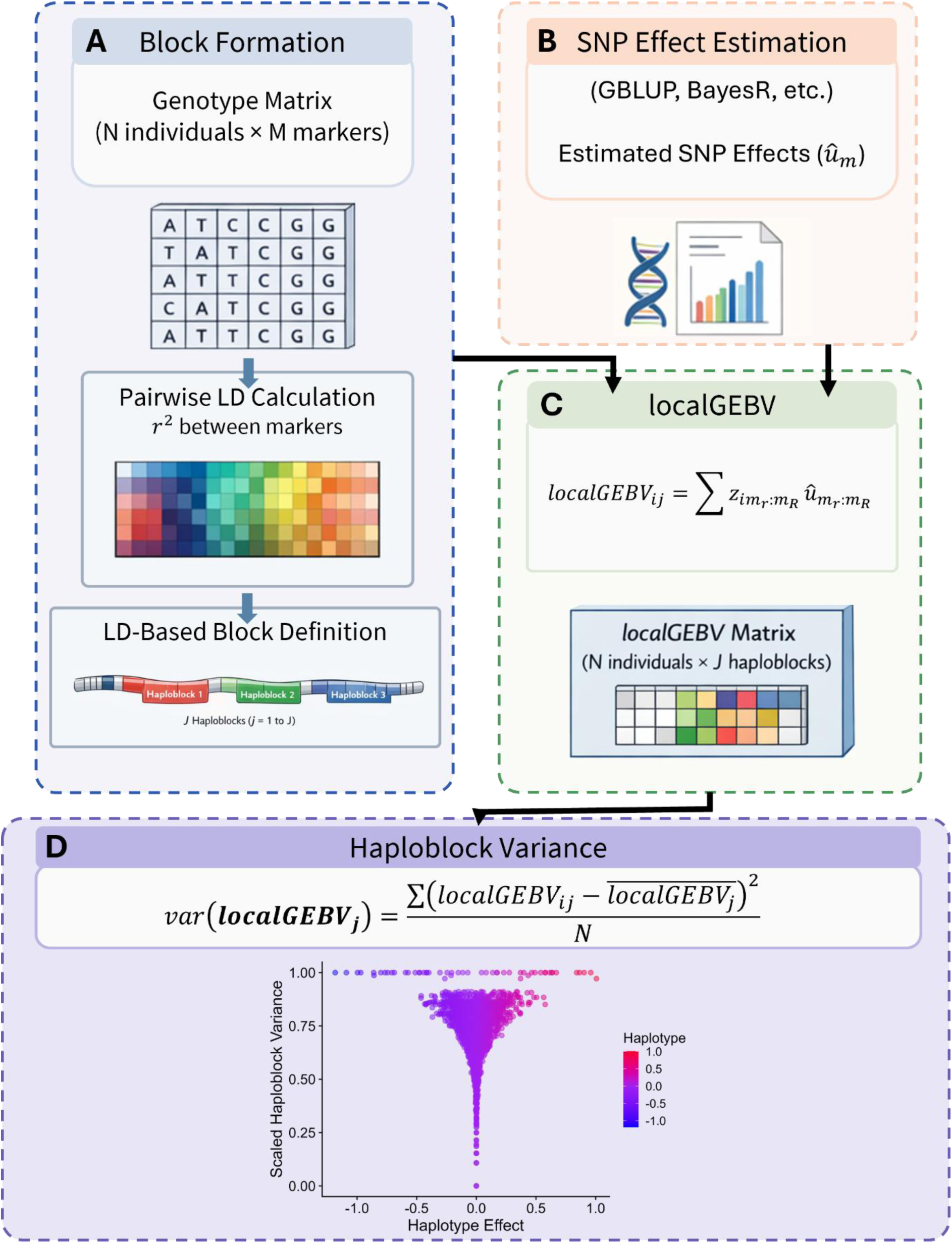
Diagram of the localGEBV process. The localGEBV process consists of three computing steps and a summarization step: (**A**) block formation from pairwise LD between 𝑀 markers, (**B**) SNP effect estimation, (**C**) calculating localGEBV of all 𝑁 individuals (denoted by 𝑖) for all 𝑗 individual haploblocks in 𝐽, and (**D**) computing the variance of all 𝑗 individual haploblocks in 𝐽 (tornado plot is for demonstration purposes only).

